# New, biomechanically sound tendon tissue after injection of uncultured, autologous, adipose derived regenerative cells in partial Achilles tendon defects in rabbits

**DOI:** 10.1101/2024.02.18.580890

**Authors:** Christoph Schmitz, Christopher Alt, Tobias Würfel, Stefan Milz, Jacqueline Dinzey, Ashley Hill, Katie J. Sikes, Lindsey Burton, Jeremiah Easley, Holly L. Stewart, Christian M. Puttlitz, Benjamin C. Gadomski, Kevin Labus, David A. Pearce, Nicola Maffulli, Eckhard U. Alt

**Author notes:** Correspondence Chair of Anatomy II, Institute of Anatomy, Faculty of Medicine, Ludwig-Maximilians University, Pettenkoferstr. 11, 80336 Munich, Germany. This paper contains data from the MD thesis of Tobias Würfel.

## Abstract

**Background:** Current management options for partial tendon tears may not offer future potential to heal tissue and improve clinical results. This study tested the hypothesis that treatment of a partial rabbit common calcaneus tendon (CCT) defect with uncultured, autologous, adipose derived regenerative cells (UA-ADRCs) enables regenerative healing without scar formation, as recently observed in a biopsy of a human supraspinatus tendon.

**Methods:** A full-thickness hole (diameter, 3 mm) was punched into the midsubstance of the right gastrocnemius tendon (GT; which is a part of the CCT) of adult, female New Zealand white rabbits. Immediately thereafter the rabbits were treated by application of an averaged 28.3×10^6^ UA-ADRCs in 0.5 ml lactated Ringer’s solution (RLS) into the GT defect and surrounding tendon tissue, or underwent sham treatment. Rabbits were sacrificed either four weeks (W4) or twelve weeks (W12) post-treatment, and the CCTs were investigated using histology, immunohistochemistry and non-destructive biomechanical testing.

**Results:** Newly formed connective tissue was consistent with the formation of new tendon tissue after treatment with UA-ADRCs, and with the formation of scar tissue after sham treatment, at both W4 and W12 post-treatment. Biomechanical testing demonstrated a significantly higher mean percent relaxation after treatment with UA-ADRCs than after sham treatment (p < 0.05), and significant, negative correlations between the peak stress as well as the equilibrium stress and the cross-sectional area of the CCT (p < 0.05) after treatment with UA-ADRCs but not after sham treatment.

**Conclusions:** Management of partial tendon tears with UA-ADRCs has the potential to be truly “structure-modifying”.

## INTRODUCTION

Current clinical treatment options for tendon tears offer only limited potential for true tissue healing and improvement of clinical results. Treatment of symptomatic, partial-thickness rotator cuff tears (sPTRCT) with uncultured, unmodified, autologous, adipose derived regenerative cells (UA-ADRCs) isolated from lipoaspirate at the point of care is safe and more effective than corticosteroid injection [1,2]. To this end, subjects aged between 30 and 75 years with sPTRCT who did not respond to physical therapy treatments for at least six weeks were randomly assigned to receive either a single injection of an average 11.4×10^6^ UA-ADRCs (in 5 mL liquid; mean cell viability: 88%) or a single injection of 80 mg of methylprednisolone (40 mg/ml; 2 mL) plus 3 mL of 0.25% bupivacaine. The UA-ADRCs were isolated from autologous lipoaspirate at the point of care using the Transpose RT system (InGeneron Inc., Houston, TX, USA) [3-5]. No severe adverse events related to the injection of UA-ADRCs were observed during the follow-up period. The risks connected with treatment of sPTRCT with injection of UA-ADRCs were not greater than those connected with treatment of sPTRCT with injection of corticosteroid. Compared with injection of corticosteroid, injection of UA-ADRCs resulted in significantly higher mean ASES Total scores at W24, W52 and M41, a significantly higher mean SF-36 Total score at W24, and significantly higher mean VAS Pain scores at W24 and W52 post-treatment (p<0.05). The use of UA-ADRCs in subjects with sPTRCT is safe and leads to improved shoulder function without adverse effects. To verify the results of these studies in a larger patient population, a randomized controlled trial on 246 patients suffering from sPTRCT is currently ongoing [6].

In animal models, injections of adult stem cells isolated from adipose tissue into pathologic tendon tissue has produced a positive biological response [7-10]. Reported beneficial effects include a decreased number of inflammatory cells, improved regeneration of tendons with less scarred healing, improved collagen fiber arrangement, higher load-to-failure, and higher tensile strength of the treated tendons [7-10]. However, it has remained unknown whether such beneficial effects could also be observed in a human tendon treated with UA-ADRCs. Therefore, we recently performed, for the first time, a comprehensive histological and immunohistochemical analysis of the biopsy of a supraspinatus tendon of a then 66-year-old patient with traumatic rotator cuff injury, taken ten weeks after local injection of UA-ADRCs [11]. The UA-ADRCs were isolated from autologous lipoaspirate at the point of care using the same system as used in [1-6]. Our analysis demonstrated clear evidence towards regenerative healing of the injured supraspinatus tendon [11]. Of note, no formation of adipocytes was observed. These findings indicate that injected UA-ADRCs can indeed form new tendon tissue and regenerate an injured human tendon.

This study tested the following hypothesis using a rabbit common calcaneal tendon (CCT) defect model: treatment of a partial CCT defect (induced by punching a full-thickness hole (diameter, 3 mm) into the midsubstance of the right gastrocnemius tendon, which is a part of the CCT) by treatment with UA-ADRCs isolated from autologous adipose tissue [1-6,11] results in faster and better tendon regeneration than by application of Ringer’s lactate solution (RLS), and histological and immunohistochemical findings obtained on this animal model are comparable with the results reported in [11]. Furthermore, we hypothesized that the biomechanical properties of the CCT were better restored after treatment with UA-ADRCs than after sham treatment.

## METHODS

### Ethics

This study was approved by the Colorado State University (CSU) Institutional Animal Care and Use Committee (Fort Collins, CO, USA) (Protocol # 1473; approval issued on February 1st, 2021 and renewed/amended on February 1st, 2022).

### Animals

A total of n=32 female New Zealand white rabbits (Oryctolagus cuniculus) were used (age range, 9 – 13 months; skeletally mature; body weight, 4.8 ± 0.6 kg (mean ± SD); range, 4.1 kg – 6.2 kg). Rabbits were obtained from Western Oregon Rabbit Co. (Philomath, OR, USA) and were housed and maintained in a temperature and humidity-controlled room at approximately 25° C with 12 hr light/dark cycles at the CSU Laboratory Animal Resources (LAR) Building (Fort Collins, CO, USA) for the entirety of the study period in accordance with LAR standard operation procedures. Rabbits were allowed to acclimatize for at least 14 days. They were single housed in standard rabbit cages on a smooth slatted floor, with waste pans below to catch feces/urine. Cages were changed out every two weeks and pan liners were changed daily. Rabbits were fed with commercial laboratory rabbit chow (Envigo Teklad 2031; Envigo, Indianapolis, IN, USA) and grass hay mix ad libitum, and were provided tap water ad libitum.

All rabbits included in this study were naïve and had never been used for any previous testing. Prior to the start of this study, rabbits underwent a physical examination by a veterinarian. All rabbits included in this study were determined to have an acceptable health status, age and weight, and were deemed free of any disease or condition that would have interfered with the purpose or conduct of this study. The body weight of the rabbits was determined bi-weekly during the entire duration of this study.

### Randomization and blinding

The first cohort of animals consisted of n=16 rabbits that were randomly assigned to four experimental groups (Groups 1-4; n=4 rabbits per group) by pull of a hat (Table 1). The same was done with the second cohort of rabbits, but the n=16 animals were randomly assigned to only two groups (Groups 5-6; n=8 rabbits per group) (Table 1).

**Table 1.** Groups of rabbits investigated in the present study.

| Group | n | Treatment | Type | Time post-treatment |
| --- | --- | --- | --- | --- |
| 1 | 4 | UA-ADRCs | H / I | W4 |
| 2 | 4 | UA-ADRCs | H / I | W12 |
| 3 | 4 | LRS | H / I | W4 |
| 4 | 4 | LRS | H / I | W12 |
| 5 | 8 | UA-ADRCs | B / I | W12 |
| 6 | 8 | LRS | B / I | W12 |

Abbreviations: n, number of animals; UA-ADRCs, uncultured, unmodified, autologous, adipose derived regenerative cells; RLS, Ringer's lactate solution; H, histology; I immunohistochemistry; B, non-destructive biomechanical analysis; W4, four weeks; W12, twelve weeks.

Experimenters and researchers were not blinded to the treatment administered to the rabbits. The histopathology slides were examined in a blinded fashion.

### Anesthesia

To harvest adipose tissue, all rabbits were placed in sternal recumbency on a surgical table. For punching a hole into the midsubstance of the right GT, rabbits in all groups were placed in left lateral recumbency on a surgical table.

Rabbits were maintained and monitored (approximately every five minutes) under general anesthesia for the surgical procedure. The anesthetic protocol comprised preoperative subcutaneous injection of Buprenorphine (0.03 mg/kg bodyweight (BW) (Par Pharmaceutical, Chestnut Ridge, NY, USA), Glycopyrrolate (0.005 mg/kg BW) (Somerset Therapeutics, Hollywood, FL, USA), Ketamine (25 mg/kg BW) (Dechra Veterinary Products, Overland Park, KS, USA) and Dexmedetomidine (0.02 mg/kg BW) (Dechra Veterinary Products), and inhalation of Isoflurane (1-5% continuous during surgery) (VetOne, Boise, ID, USA). Monitoring focused on heart rate, respiratory rate, ECG, pulse oximetry, CO2 exhalation, and reflexes.

### Harvesting of adipose tissue and isolation of UA-ADRCs

The interscapular region of each rabbit in Groups 1, 2 and 5 was shaved and prepped for aseptic surgery using alternating scrubs of povidone-iodine and alcohol. Then, a 2 cm long incision was made over the interscapular fat depot. The fat pad was dissected carefully and collected as previously described [12]. On average, 21.1 ± 5.3 (mean ± SD) grams of adipose tissue (median, 21.5 g; range, 11.6 g – 30.5 g) were collected. Adipose tissue was stored in a sterile tube in sterile RLS and processed immediately. The skin was then closed routinely with nonabsorbable nylon monofilament suture (Covidien, Mansfield, MA, USA).

Following collection of adipose tissue, UA-ADRCs were isolated using the Transpose RT system (InGeneron) as described in [1,3] and characterized prior to implantation. The mean total cell yield was 14.5×10^5^ ± 7.1×10^5^ cells / g adipose tissue (median, 13.2×10^5^ cells / g adipose tissue; range, 4.7×10^5^ – 29.5×10^5^ cells / g adipose tissue), the mean cell viability was 82.6% ± 5.4% (median, 83.1%; range, 75.0% – 89.9%), and the average live cell yield was 11.8×10^5^ ± 5.3×10^5^ cells / g adipose tissue (median, 10.8×10^5^ cells / g adipose tissue; range, 3.6×10^5^ – 22.3×10^5^ cells / g adipose tissue).

All processing was conducted under sterile conditions, and cells were maintained sterilely until implantation. Each batch of fat (from individual rabbits) was processed separately to avoid contamination between batches and was given a unique identification/lot number for reference.

### Experimental procedure

The right hindlimb of all rabbits was shaved and prepped for aseptic surgery using alternating scrubs of povidone-iodine and alcohol. Then, the right CCT was approached from the posterior side using a small incision (1-3 cm) for visualization of the target area. The peritenon was dissected away from the CCT, and the GT was dissected away from the superficial digital flexor tendon (c.f. [13]). A full thickness defect (diameter, 3 mm) was made with a punch in the midsubstance of the GT approximately 2.5 cm from the calcaneus insertion as shown in Figure 1 (c.f. [14]).

**Fig. 1.**
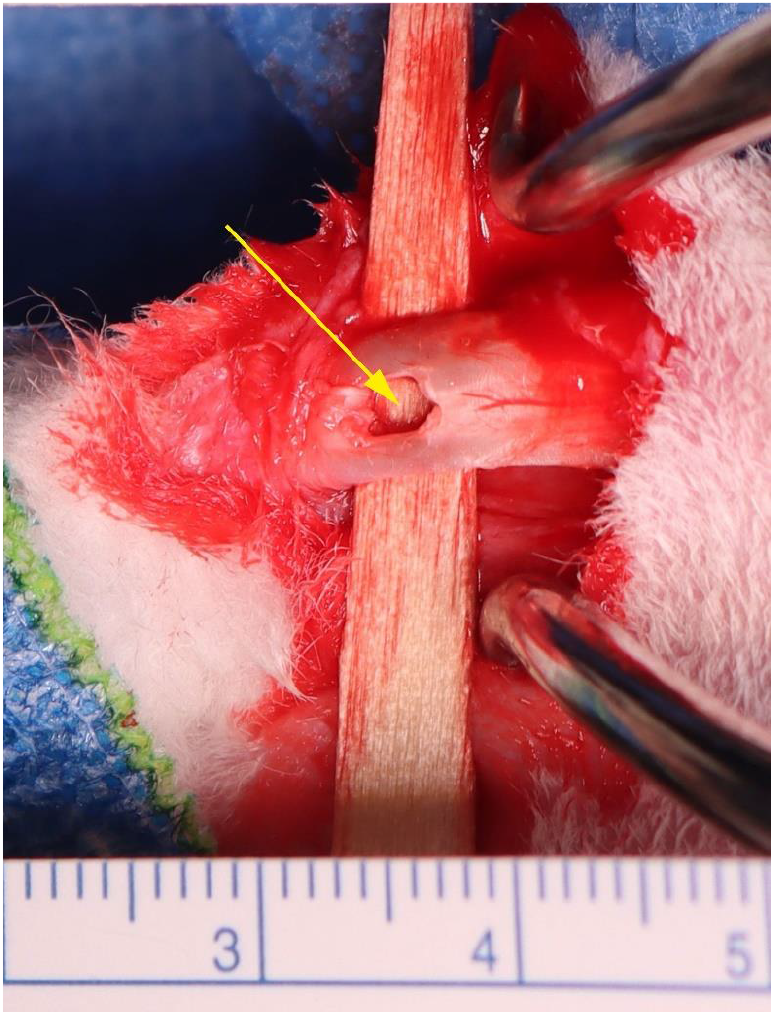
Production of a full thickness defect in the midsubstance of the rabbit gastrocnemius tendon. The defect (yellow arrow) was produced with a punch with a diameter of 3 mm, approximately 2.5 cm from the calcaneus insertion.

A surgical ruler was used to determine the distance of the defect from the calcaneus to assist with post-mortem analysis. The peritenon was partially closed with 4-0 monofilament absorbable sutures (Biosyn; Medtronics, Minneapolis, MN, USA). Then, rabbits in Groups 1, 2 and 5 received an injection of on average 28.3×10^6^ ± 11.6×10^6^ UA-ADRCs (mean ± SD) (median, 24.5×10^6^; range, 10.9×10^6^ – 58.5×10^6^) in 0.5 ml RLS (rabbits in Groups 1 and 2) or 0.5 ml saline (rabbits in Group 5) into the defect and surrounding tendon tissue. Rabbits in Groups 3 and 4 received an injection of 0.5 ml RLS into the defect and surrounding tendon tissue, and rabbits in Group 6 an injection of 0.5 ml saline. Following injection, the remainder of the peritenon was closed and the skin was then closed routinely with nonabsorbable nylon monofilament suture (Covidien). The left CCT was left intact.

Buprenorphine (0.03 mg/kg BW; Par Pharmaceutical) was injected subcutaneously 12- and 24-hours post-surgery, and Meloxicam (1 mg/kg BW; VetOne) was injected subcutaneously every 24 hours post-op up to 7 days post-op.

### Euthanasia

Once initially sedated using Ketamine (5-30 mg/kg BW) (Dechra Veterinary Prod-ucts), and Dexamedetomidine (0.05-0.125 mg/kg BW) (Dechra Veterinary Products) or Xylazine (5 mg/kg BW) (VetOne), rabbits were placed on isoflurane face mask at 4-5% until they reached a surgical plane of anesthesia, then maintained on 2-3% isoflurane. All rabbits were humanely euthanized by intravenous overdose of Pentobarbitone sodium (88 mg/kg BW) (Dechra Veterinary Products), in accordance with the American Veterinary Medical Association guidelines. The survival time points of rabbits in Groups 1 and 3 were four weeks (+/-4 days) post-surgery/treatment or sham treatment, and twelve weeks (+/-4 days) post-surgery/treatment or sham treatment for rabbits in Groups 2, 4, 5 and 6.

### Preparation of histological sections

Both hind limbs were collected en bloc from each rabbit. The limbs were carefully dissected of extraneous muscle and soft tissue to isolate the CCT and associated treatment region. For Groups 1-4, tendon samples were then transected from the gastrocnemius muscle and calcaneus bone and were further sectioned by creating a block of tissue approximately 3 cm long around the injection site. For Groups 5 and 6, the limbs were dissected to isolate the CCT, associated treatment region, and the calcaneal attachment. Following dissection, all samples were placed in 10% neutral buffered formalin (NBF) until fixation was complete. Samples from Groups 5 and 6 underwent decalcification using 5% formic acid, and the decalcification endpoint was determined radiographically. Decalcification was not performed on Groups 1-4.

All samples were processed using standard paraffin techniques and longitudinally sectioned at 5 μm thickness; sections were mounted on positive charged glass slides. Three sections were cut per limb and stained with respectively Azan Trichrome, Picrosirius Red, Hematoxylin and Eosin (H&E), or Safranin O/Fast Green and cover slipped using DPX mountant (Sigma Aldrich, St. Louis, MO, USA). Additional sections were cut but not stained. Stained and unstained sections were shipped to the laboratory of the Department of Anatomy II at LMU Munich (Munich, Germany) (LMU-Anatomy).

### Biomechanical analysis

Both hind limbs of rabbits in Groups 5-6 were collected en bloc from each rabbit. The limbs were carefully dissected of extraneous muscle and soft tissue to isolate the CCT and associated treatment region. Then, tendons with remnants of the adjacent muscles (lateral head of the gastrocnemius, medial head of the gastrocnemius and superficial digital flexor) and the calcaneus bone were transected from the other tissue.

Following dissection, the length and cross sectional area (assuming rectangular geometry) of the tendon specimens was measured using calipers without applying any deforming force to the tendons.

A non-destructive biomechanical analysis was performed to quantify changes in the mechanical characteristics of the CCT / calcaneus constructs under physiological loads. Sample hydration was maintained via physiologic saline spray at approximately ten-minute intervals during the entire preparation and testing protocol. The calcaneus was rigidly clamped to the frame of a servo-hydraulic material testing system (MTS) (MiniBionix 858; MTS, Eden Prairie, MN, USA), such that the CCT physiological loading direction was aligned axially with the actuator of the MTS as shown in Figure 2. At its proximal end the tendon was gripped in a clamp rigidly attached to a force transducer and the actuator.

**Fig. 2.**
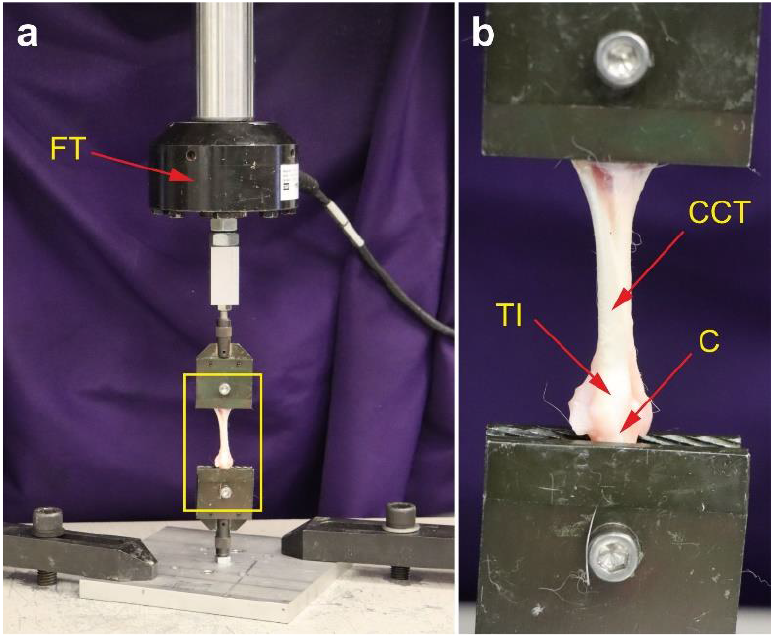
Non-destructive biomechanical analysis of a rabbit common calcaneal tendon ex vivo using a servo-hydraulic material testing system (MiniBionix 858; MTS, Eden Prairie, MN, USA). The yellow rectangle in (a) indicates the position of the detail shown in (b). Abbreviations: FT, force transducer; CCT, common calcaneal tendon; TI, tendon insertion; C, calcaneus.

The biomechanical testing procedure consisted of an initial 2 N static pre-load maintained for two minutes, followed by preconditioning cycles and a stress-relaxation test. Following the pre-load, the reference gauge length of the tendon was measured and recorded as the distance between gripping surfaces in mm. The measured gauge length was used to determine the applied displacement for preconditioning cycles and stress-relaxation testing. Afterwards the samples underwent 10 cycles of preconditioning between 0% and 2% engineering strain. Following preconditioning, specimens were loaded in one stress-relaxation test. A magnitude of 5% engineering strain was applied to the specimens at a rate of 5 mm/s. The 5% strain was maintained constant for a 100 s relaxation period. Force and displacement data were collected throughout the stress-relaxation test.

The following structural properties representing the mechanical behavior of the construct were calculated from the force and displacement data collected during the stress-relaxation test: Peak Load (N) (the maximum force during the test) and Equilibrium Load (N) (the force at the end of the 100 s relaxation period).

Furthermore, the following material properties were calculated by normalization of the structural properties to the cross-sectional area: Peak Stress (MPa) (the maximum stress during the test), Equilibrium Stress (N) (the stress at the end of the 100 s relaxation period) and Percent Relaxation (%) (the percent difference between the peak stress and equilibrium stress).

### Immunohistochemistry

Immunohistochemistry was performed at LMU-Anatomy on de-paraffinized and rehydrated sections that were washed with phosphate buffered saline (PBS) containing Tween 20 (Sigma Aldrich, St. Louis, MO, United States). After antigen retrieval and / or enzymatic and / or chemical pretreatment the slides were blocked with different solutions for 15 min to 60 min at room temperature (details are provided in Table 2). Then, sections were incubated with primary antibodies for the detection of procollagen 1, type III collagen, CD163 and aggrecan as summarized in Table 2.

**Table 2.** Characteristics of the antibodies used in the present study.

|  |  |  |
| --- | --- | --- |
| Procollagen 1 | Immunoglobuline isotype / clone status<br>Catalog no. / provider<br>Demasking of antigen<br>Blocking<br>Dilution and incubation parameters<br>Secondary antibody used | IgG1 / mouse, monoclonal<br>SP1.D8 / DSHB<sup>a</sup><br>Not applicable<br>Vector Bloxall SP-6000<sup>b</sup>, LOT ZG1216 from April 04, 2021, and normal horse serum blocking solution 2.5% (S-2012-50)<sup>b</sup><br>1:10, 4 °C, over night<br>Horse-anti-mouse IgG BA-2000<sup>b</sup>, 1:200 |
| Type III collagen | Immunoglobuline isotype / clone status<br>Catalog no. / provider<br>Demasking of antigen<br>Blocking<br>Dilution and incubation parameters<br>Secondary antibody used | IgG1 / mouse, monoclonal<br>C7805 (Clone FH-7A) / Sigma-Aldrich (St. Louis, MO, United States)<br>Protease XIV<br>Vector Bloxall SP-6000<sup>b</sup>, LOT ZG1216 from April 04, 2021, and normal horse serum blocking solution 2.5% (S-2012-50)<sup>b</sup><br>1:150, 4 °C, over night<br>Horse-anti-mouse IgG BA-2000<sup>b</sup>, 1:200 |
| CD163 | Immunoglobuline isotype / clone status<br>Catalog no. / provider<br>Demasking of antigen<br>Blocking<br>Dilution and incubation parameters<br>Secondary antibody used | IgG 1/ mouse, monoclonal<br>5C6 FAT BMA Biomedicals (Augst, Switzerland)<br>Not applicable<br>3% H<sub>2</sub>O<sub>2</sub> in Methanol<br>1:400, 4°C, over night<br>Horse-anti-mouse IgG BA-2000<sup>b</sup>, 1:200 |
| Aggrecan | Immunoglobuline isotype / clone status<br>Catalog no. / provider<br>Demasking of antigen<br>Blocking<br>Dilution and incubation parameters<br>Secondary antibody used | IgG1 / mouse, monoclonal<br>12/21/1-C-6 / DSHB<sup>a</sup><br>3% H<sub>2</sub>O<sub>2</sub> in Methanol / Chondroitinase AC<sup>3</sup><br>Normal horse serum blocking solution 2.5% (S-2012-50)<sup>2</sup><br>1:5, 4 °C, over night<br>Horse-anti-mouse IgG BA 2000<sup>b</sup>, 1:200 |

<sup>a</sup>Antibodies SP1.D8 (developed by Dr. H. Furthmayr) and 12/21/1-C-6 (developed by Dr. B. Caterson) were obtained from the Developmental Studies Hybridoma Bank (DSHB), created by the Eunice Kennedy Shriver National Institute of Child Health and Human Development of the National Institutes of Health of the United States, and maintained at The University of Iowa, Department of Biology, Iowa City, IA, United States. <sup>2</sup>Provider: Vector Laboratories (Burlingame, CA, United States). <sup>3</sup>The 12/21/1-C-6 antibody recognizes an epitope on the core protein of the aggrecan molecule. Therefore, the pre-treatment requires the removal of the disaccharide side chains and the removal of oxidation effects [15].

Antibody binding was detected with the Vectastain Elite ABC Kit Peroxidase (HRP) (Vector Laboratories, Burlingame, CA, United States) with a secondary antibody incubation of 30 min for all slides (Table 2). Visualization of peroxidase activity was performed using diaminobenzidine (Vector Impact DAB chromogen solution; Vector Laboratories), resulting in a brown staining product. The sections were counterstained with Mayer’s hematoxylin. Primary antibodies were omitted and replaced with PBS in order to perform specificity controls. Microscopic evaluations were performed by C.S., C.A., S.M. and E.A.

### Design-based stereologic analysis

Relative amounts (area/area) of cells, vessels and extracellular matrix in newly formed connective tissue were determined on sections stained with hematoxylin and eosin of the right CCTs using point counting as described in detail in [16]. To this end, newly formed connective tissue adjacent to original tendon tissue was delineated, and individual point counting grids were created that resulted in a mean number of 501 (range, 490-516) points per section. Analyses were performed with a computerized stereology workstation, consisting of a modified light microscope (Axioskop; Carl Zeiss Microscopy, Jena, Germany) with Plan-Neofluar objectives 1.25× (numerical aperture [NA] = 0.03), 2.5× (NA = 0.085), Plan-Apochromat objectives 5× (NA = 0.16), 10× (NA = 0.45), 20× (NA = 0.8) and 40× (NA = 0.95) (Carl Zeiss Microscopy), motorized specimen stage (MBF Bioscience, Williston, VT, USA), stage controller (MAC 6000 XY; Ludl Electronics) focus encoder (MT 1271; Heidenhain, Traunreut, Germany), CCD color video camera (1,600 × 1,200 pixels; MBF Bioscience, Williston, VT, USA), and stereology software (Stereo Investigator Version 11.01.2 64 bit; MBF Bioscience).

### Determination of Bonar scores

The Bonar scores cell morphology, cellularity, vascularity, and ground substance were determined for newly formed connective tissue according to [17-19] on sections of the right CCT stained with hematoxylin and eosin. The Bonar score collagen arrangement was determined for newly formed connective tissue on sections of the right CCT stained with Picrosirius Red and evaluated using polarization microscopy.

### Statistical analysis

The histological findings between Groups 5 and 6 shown in Figures 14 and 16 were compared using Fisher’s exact test.

Mean and standard deviation were calculated from the results of the design-based stereologic analysis, tendon measurements and biomechanical analysis.

The results of the design-based stereologic analysis were compared between Groups 1-4 using two-way ANOVA (two treatments; two time points), followed by Bonferroni’s multiple comparisons test (UA-ADRCs vs RLS).

Bonar scores were compared between Groups 1 and 3 as well as between Groups 2 and 4. Comparisons were performed using the Mann Whitney test.

The results of the biomechanical analysis were compared between the left (intact / untreated) and the right (injured / treated or sham treated) CCTs of the rabbits in Groups 5 and 6 (i.e. four groups) using one-way ANOVA, followed by Bonferroni’s multiple comparisons test.

Correlations between the results of the design-based stereologic analysis and the number of injected cells, as well as between the results of the biomechanical analysis and the number of injected cells, were tested using linear regression analysis.

In all analyses, an effect was considered statistically significant if its associated p value was smaller than 0.05. Calculations were performed using GraphPad Prism (Version 10.1.2 for Windows; GraphPad Software, San Diego, CA, USA).

### Photography

All photomicrographs were produced by digital photography.

The photomicrographs shown in Figures 1-4, 7-10, 11e-h and m-p, 12e-h and m-p, and 13-18 were created from virtual slides (using the software Biolucida Viewer; Version 2020.1.0; MBF Bioscience) that were produced by digital photography using an automated scanning microscopy workstation. The latter consisted of a M2 AxioImager microscope (Zeiss, Goettingen, Germany), 10× Plan-Apochromate objective (NA = 0.3; Zeiss), 2-axis computer-controlled stepping motor system (4”× 3” XY; Prior Scientific, Jena, Germany), focus encoder (Heidenhain, Traunreut, Germany) and color digital camera (AxioCam MRc; 2/3” CCD sensor, 1388 × 1040 pixels; Zeiss). The whole system was controlled by the software Stereo Investigator (Version 11.06.2; MBF Bioscience). On average 670 (range, 211-953) images were captured for each composite. These images were made into one montage each using the Virtual Slide module of the Stereo Investigator software (MBF Bioscience); the size of the resulting 2D virtual slides varied between 67 MB and 359 MB. The photomicrographs shown in Figures 5, 6, 11a-d and i-l, and 12a-d and i-l were produced using a Zeiss Axiophot Microscope equipped with an Axiocam HRc digital camera (2/3” CCD sensor, 1388 × 1040 pixels; Zeiss) that was controlled by the software Zeiss Axiovision SE64 (Rel. 4.9.1 SP2). The images were taken in transmitted light mode either without (Fig. 5a-d and i-l, and Fig. 6a-d and i-l) or with polarized light (Fig. 5e-h and m-p, Fig. 6e-h and m-p, Fig. 11a-d and i-l, and Fig. 12a-d and i-l) using a 5× Zeiss Plan-Neofluar objective (NA = 0.15). The polarized images were taken in black and white mode of the digital camera. Illumination was adjusted using the automatic measurement function of the Zeiss Axiovision software.

**Fig. 3.**
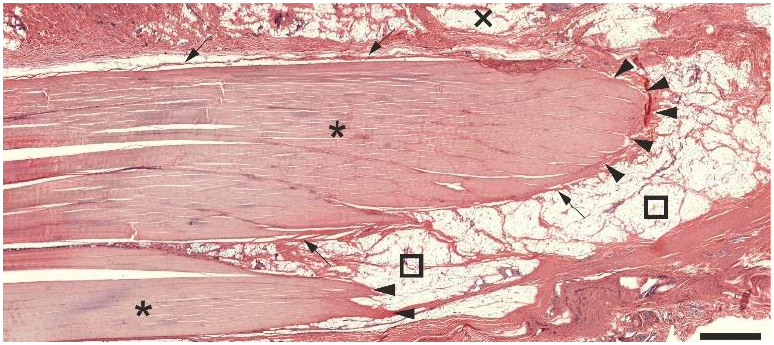
Low-power photomicrograph of a representative 5 μm thick section stained with hematoxylin and eosin of a left common calcaneal tendon of a rabbit (no surgery, no treatment). Details are in the text. The scale bar represents 1 mm.

**Fig. 4.**
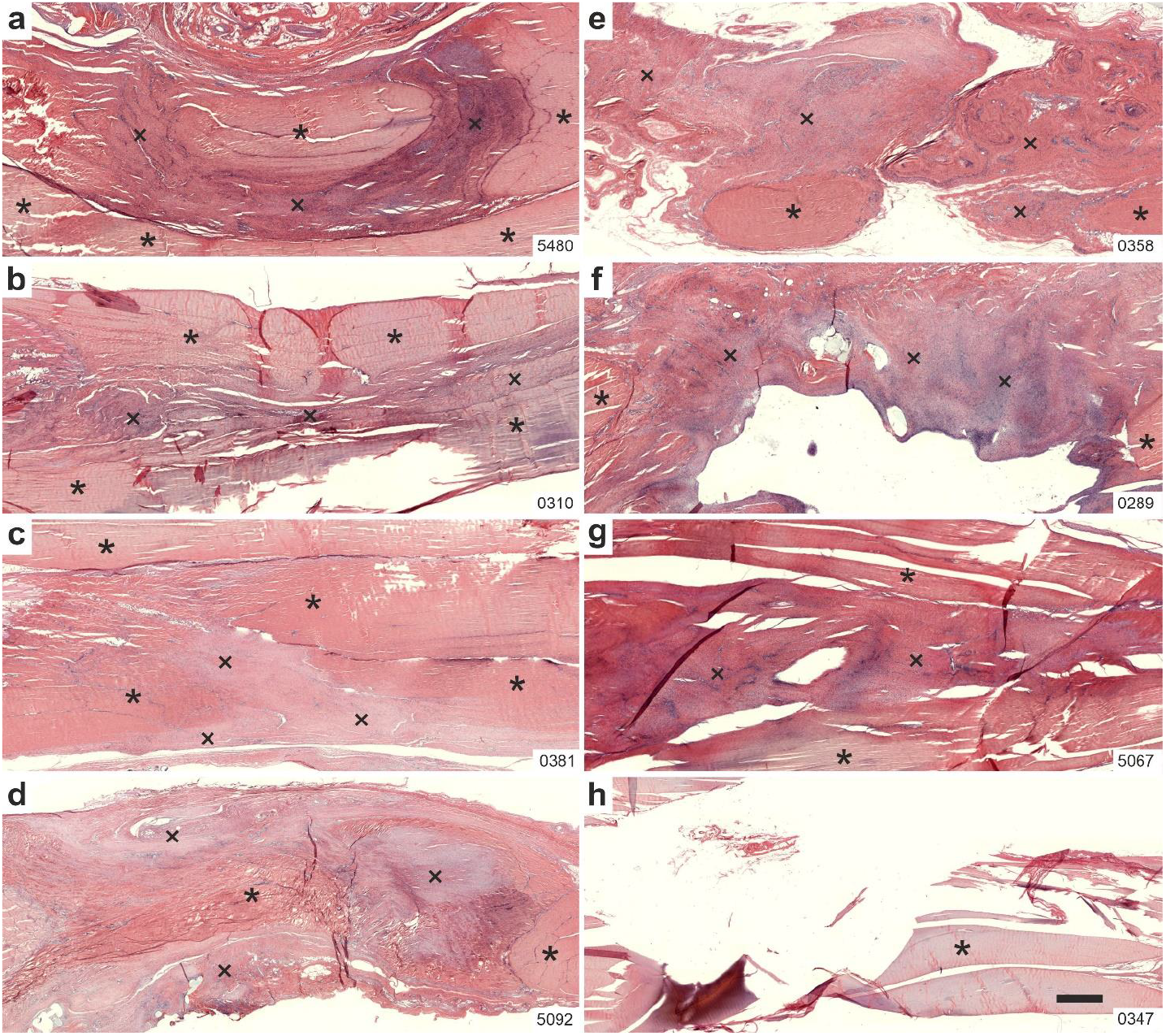
Representative low-power photomicrographs of 5 μm thick sections stained with hematoxylin and eosin of the right (surgery, treatment or sham treatment) common calcaneal tendon of the rabbits in Group 1 (UA-ADRCs / W4) (a-d) and in Group 3 (RLS / W4) (e-h). The asterisks indicate original tendon tissue, and the crosses point to newly formed connective tissue. The numbers in the lower right corner of each panel identify the individual animals according to the study protocol. The scale bar in h represents 1 mm in all panels. Details are in the text.

**Fig. 5.**
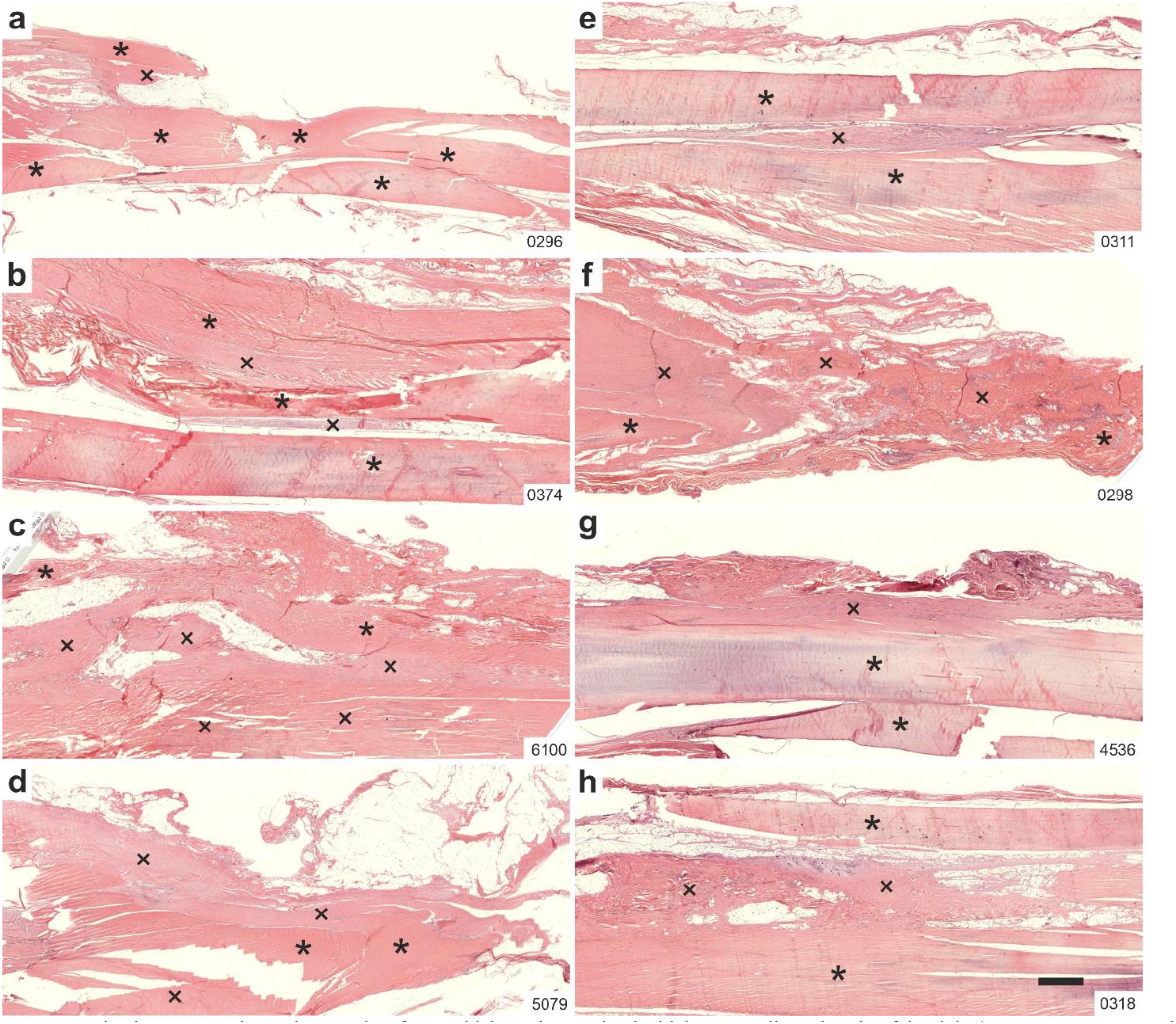
Representative low-power photomicrographs of 5 μm thick sections stained with hematoxylin and eosin of the right (surgery, treatment or sham treatment) common calcaneal tendon of the rabbits in Group 2 (UA-ADRCs / W12) (a-d) and in Group 4 (RLS / W12) (e-h). The asterisks indicate original tendon tissue; the crosses point to newly formed connective tissue. The numbers in the lower right corner of each panel identify the individual animals according to the study protocol. The scale bar in h represents 1 mm in all panels. Details are in the text.

**Fig. 6.**
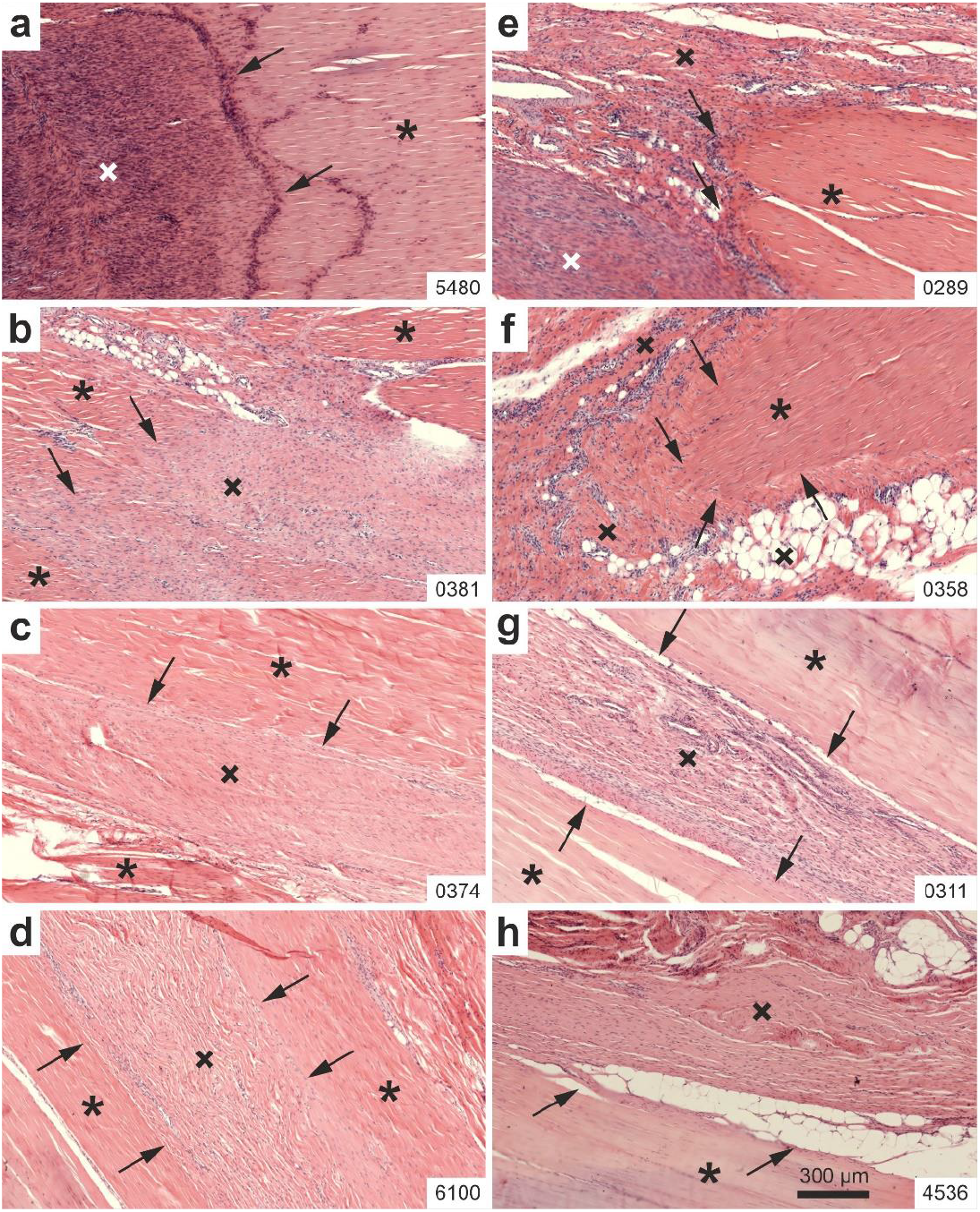
Representative high-power photomicrographs of 5 μm thick sections stained with hematoxylin and eosin of the right (surgery, treatment or sham treatment) common calcaneal tendon of two rabbits each in Group 1 (UA-ADRCs / W4) (a,b), Group 2 (UA-ADRCs / W12) (c,d), Group 3 (RLS / W4) (e,f) and Group 4 (RLS / W12) (g,h). The asterisks indicate original tendon tissue, the crosses point to newly formed connective tissue, and the arrows indicate the border between the original tendon tissue and the newly formed connective tissue. The numbers in the lower right corner of each panel identify the individual animals according to the study protocol. The scale bar in h represents 300 μm in all panels. Details are in the text.

**Fig. 7.**
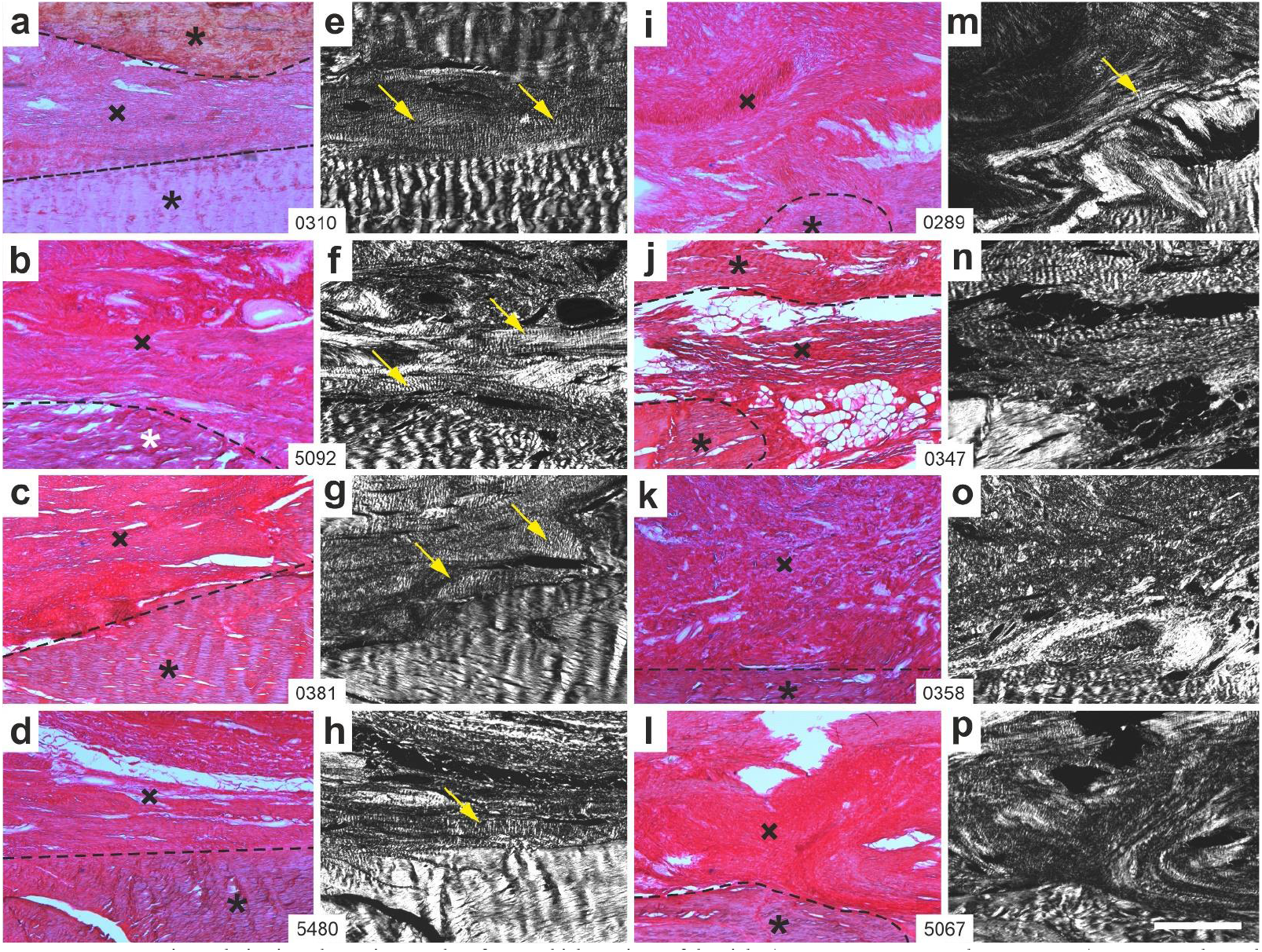
Representative polarization photomicrographs of 5 μm thick sections of the right (surgery, treatment or sham treatment) common calcaneal tendon stained with Picrosirius Red of the rabbits in Group 1 (UA-ADRCs / W4) (a-h) and in Group 3 (RLS / W4) (i-p). Panels a-d and i-l were captured using brightfield microscopy; Panels e-h and m-p show the corresponding fields of view captured using polarization microscopy. The asterisks indicate original tendon tissue, the crosses point to newly formed connective tissue, and the dashed lines indicate the border between the original tendon tissue and the newly formed connective tissue. The yellow arrows indicate organized, firm connective tissue with discernible crimp arrangement within the newly formed connective tissue. The numbers in the white boxes that connect two corresponding panels each (a/e, b/f, c/g…l/p) identify the individual animals according to the study protocol. The scale bar in p represents 500 μm in all panels. Details are in the text.

**Fig. 8.**
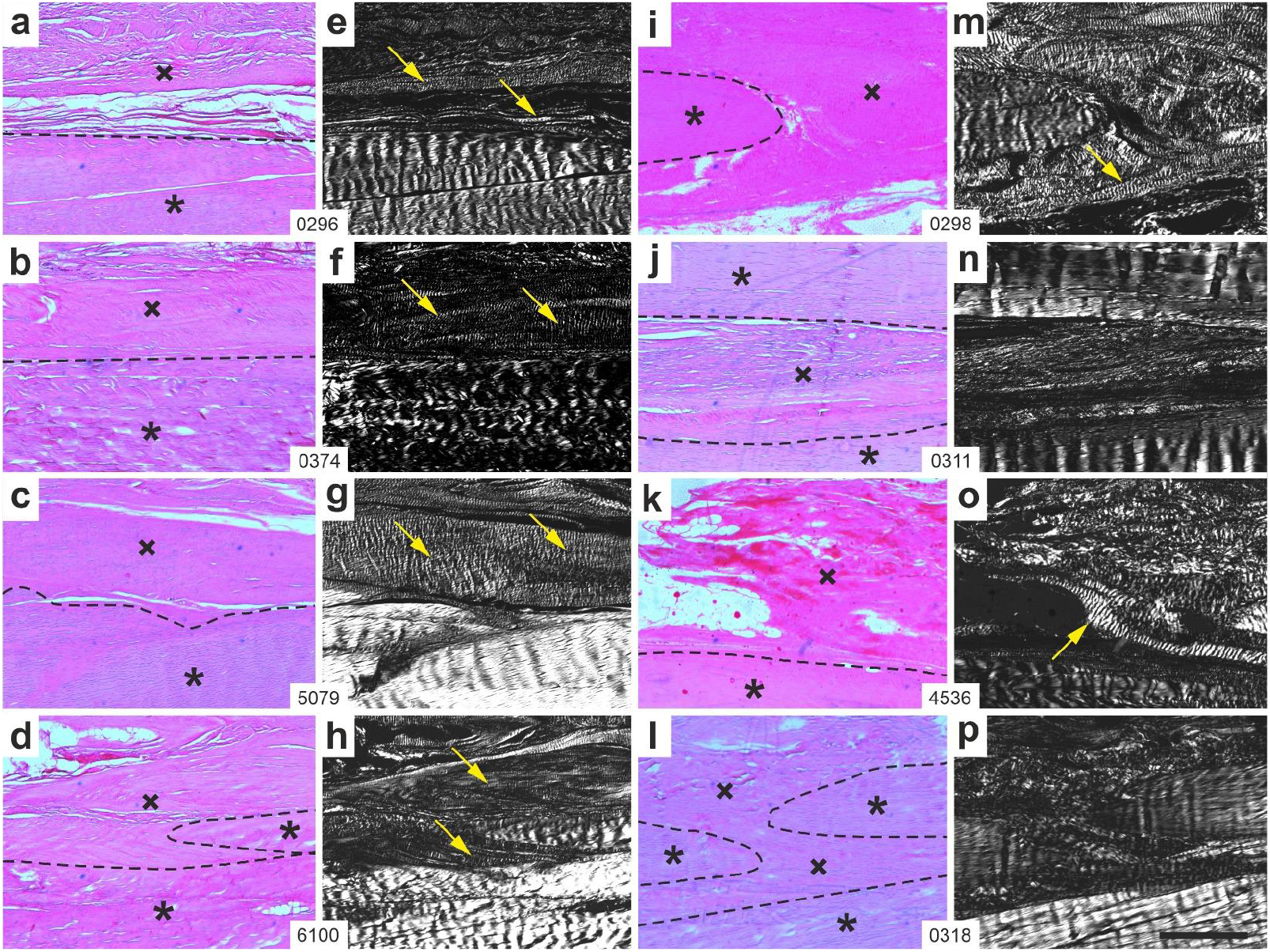
Representative polarization photomicrographs of 5 μm thick sections of the right (surgery, treatment or sham treatment) common calcaneal tendon (stained with Picrosirius Red) of the rabbits in Group 2 (UA-ADRCs / W12) (a-h) and in Group 4 (RLS / W12) (i-p). Panels a-d and i-l were captured using brightfield microscopy; Panels e-h and m-p show the corresponding fields of view captured using polarization microscopy. The asterisks indicate original tendon tissue, the crosses point to newly formed connective tissue, and the dashed lines indicate the border between the original tendon tissue and the newly formed connective tissue. The yellow arrows indicate organized, firm connective tissue with discernible crimp arrangement within the newly formed connective tissue. The numbers in the white boxes that connect two corresponding panels each (a/e, b/f, c/g…l/p) identify the individual animals according to the study protocol. The scale bar in p represents 500 μm in all panels. Details are in the text.

The final figures were constructed using Corel Photo-Paint 2021 and Corel Draw 2021 (both versions 23.1.0.389; Corel, Ottawa, Canada). Only adjustments of contrast and brightness were made using Corel Photo-Paint, without altering the appearance of the original materials.

## RESULTS

### Histology

For comparison, Figure 3 shows a representative photomicrograph of a 5 μm thick section stained with hematoxylin and eosin of a left CCT (no surgery, no treatment) of a rabbit in Group 1. The asterisks indicate tendon tissue, the arrows point to the peritenon, and the squares indicate the surrounding adipose tissue. The arrowheads point to regions where the CCT left the section plane.

Figure 4 shows representative low-power photomicrographs of 5 μm thick sections stained with hematoxylin and eosin of the right CCT of the rabbits in Group 1 (treatment with UA-ADRCs; investigation at four weeks post-treatment) (hereafter: UA-ADRCs / W4) (Fig. 4a-d) and of the rabbits in Group 3 (sham treatment; investigation at W4 post-treatment) (hereafter: RLS / W4) (Fig. 4e-h). At low magnification, the rabbits in Group 1 showed newly formed connective tissue that fully filled the gaps between the stumps of the original tendon and was homogeneously incorporated into the longitudinal structure of the tendon. In contrast, the rabbits in Group 3 showed newly formed connective tissue that did not fully fill the gaps between the stumps of the original tendon and was not homogeneously incorporated into the longitudinal structure of the tendon.

Figure 5 shows representative low-power photomicrographs of 5 μm thick sections stained with hematoxylin and eosin of the right CCT of the rabbits in Group 2 (treatment with UA-ADRCs; investigation at twelve weeks post-treatment) (hereafter: UA-ADRCs / W12) (Fig. 5a-d) and of the animals in Group 4 (sham treatment; investigation at W12 post-treatment) (hereafter: RLS / W12) (Fig. 5e-h). At low magnification, it was difficult to differentiate between the original tendon tissue and the newly formed connective tissue in the sections of the rabbits in Group 2. Both tissues were in close contact to each other. This was not the case in the sections of the rabbits in Group 4, where it was much easier to differentiate between the original tendon tissue and the newly formed connective tissue.

Figure 6 shows representative high-power photomicrographs of 5 μm thick sections stained with hematoxylin and eosin of the right CCT of two rabbits each in Group 1 (UA-ADRCs / W4) (Fig. 6a,b), Group 2 (UA-ADRCs / W12) (Fig. 6c,d), Group 3 (RLS / W4) (Fig. 6e,f) and Group 4 (RLS / W12) (Fig. 6g,h). At higher magnification, the rabbits in Groups 1 and 2 showed close contact between the original tendon tissue and the newly formed connective tissue, without the formation of blood vessels at this junction. In contrast, the animals in Group 3 showed formation of blood vessels at the junction between the original tendon tissue and the newly formed connective tissue. In addition, the rabbits in Group 4 showed formation of adipose tissue between the original tendon tissue and the newly formed connective tissue, without close contact between these tissues.

Furthermore, after treatment with UA-ADRCs (rabbits in Groups 1 and 2) the orientation of cells and extracellular matrix (ECM) in the newly formed connective tissue resembled the orientation of the cells and ECM in the original tendon tissue. This was not the case in the sections of the animals in Groups 3 and 4 (RLS). Rather, the newly formed connective tissue showed no clear orientation of cells and ECM after sham treatment.

### Polarization microscopy

Figure 7 shows representative polarization photomicrographs of 5 μm thick sections stained with Picrosirius Red of the right CCT of the rabbits in Group 1 (UA-ADRCs / W4) (Fig. 7a-h) and Group 3 (RLS / W4) (Fig. 7i-p). All rabbits in Group 1 but only one rabbit in Group 3 (Fig. 7m) showed newly formed, organized, firm connective tissue with discernible crimp arrangement at W4 post-treatment.

Figure 8 shows representative polarization photomicrographs of 5 μm thick sections stained with Picrosirius Red of the right CCT of the rabbits in Group 2 (UA-ADRCs / W12) (Fig. 8a-h) and Group 4 (RLS / W12) (Fig. 8i-p). All rabbits in Group 2 but only two rabbits in Group 4 (Fig. 8m, o) showed newly formed, organized, firm connective tissue with discernible crimp arrangement at W12 post-treatment.

### Immunolabeling of type I procollagen

Figure 9 shows representative photomicrographs of immunohistochemical detection of type I procollagen in 5 μm thick sections of the right CCT of three rabbits each in Group 1 (UA-ADRCs / W4) (Fig. 9a-c), Group 3 (RLS / W4) (Fig. 9d-f), Group 2 (UA-ADRCs / W12) (Fig. 9g-i) and Group 4 (RLS / W12) (Fig. 9j-l).

**Fig. 9.**
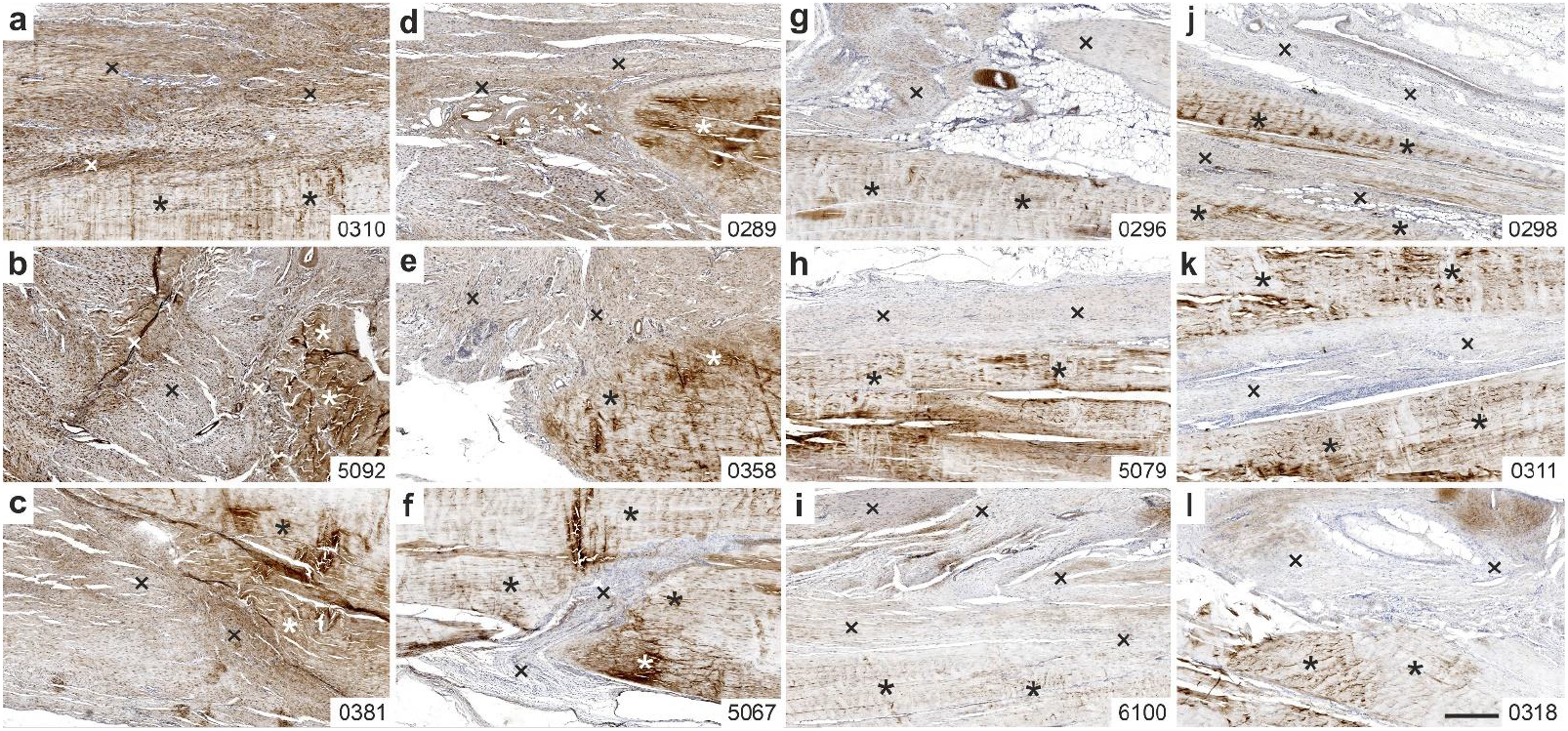
Representative photomicrographs of immunohistochemical detection of type I procollagen in 5 μm thick sections of the right (surgery, treatment or sham treatment) common calcaneal tendon of three rabbits each in Group 1 (UA-ADRCs / W4) (a-c), Group 3 (RLS / W4) (d-f), Group 2 (ADRCs / W12) (g-i) and Group 4 (RLS / W12) (j-l). The asterisks indicate original tendon tissue, and the crosses point to newly formed connective tissue. The numbers in the lower right corner of each panel identify the individual animals according to the study protocol. The scale bar in l represents 500 μm in all panels. Details are in the text.

All investigated rabbits in Group 1 (UA-ADRCs / W4) showed strong, extracellular immunolabeling for type I procollagen in newly formed connective tissue (Fig. 9a-c). Two of the three investigated rabbits in Group 3 (RLS / W4) also showed extracellular immunolabeling for type I procollagen in newly formed connective tissue, albeit more discreetly than the rabbits in Group 1 (Fig. 9d,e). In contrast, one of the three investigated rabbits in Group 3 showed almost no immunolabeling for type I procollagen in newly formed connective tissue (Fig. 9f). Furthermore, all investigated rabbits in Group 2 (UA-ADRCs / W12) showed extracellular immunolabeling for type I procollagen in newly formed connective tissue (Fig. 9g-i), albeit less pronounced than the animals in Group 1 (UA-ADCs / W4). In contrast, all investigated animals in Group 4 (RLS / W12) showed almost no immunolabeling for type I procollagen in newly formed connective tissue (Fig. 9j-l).

### Immunolabeling of type III collagen

Figure 10 shows representative photomicrographs of immunohistochemical detection of type III collagen in 5 μm thick sections of the right CCT of the rabbits in Group 1 (UA-ADRCs / W4) (Fig. 10a-d), Group 3 (RLS / W4) (Fig. 10e-h), Group 2 (UA-ADRCs / W12) (Fig. 10i-l) and Group 4 (Fig. 10m-p).

**Fig. 10.**
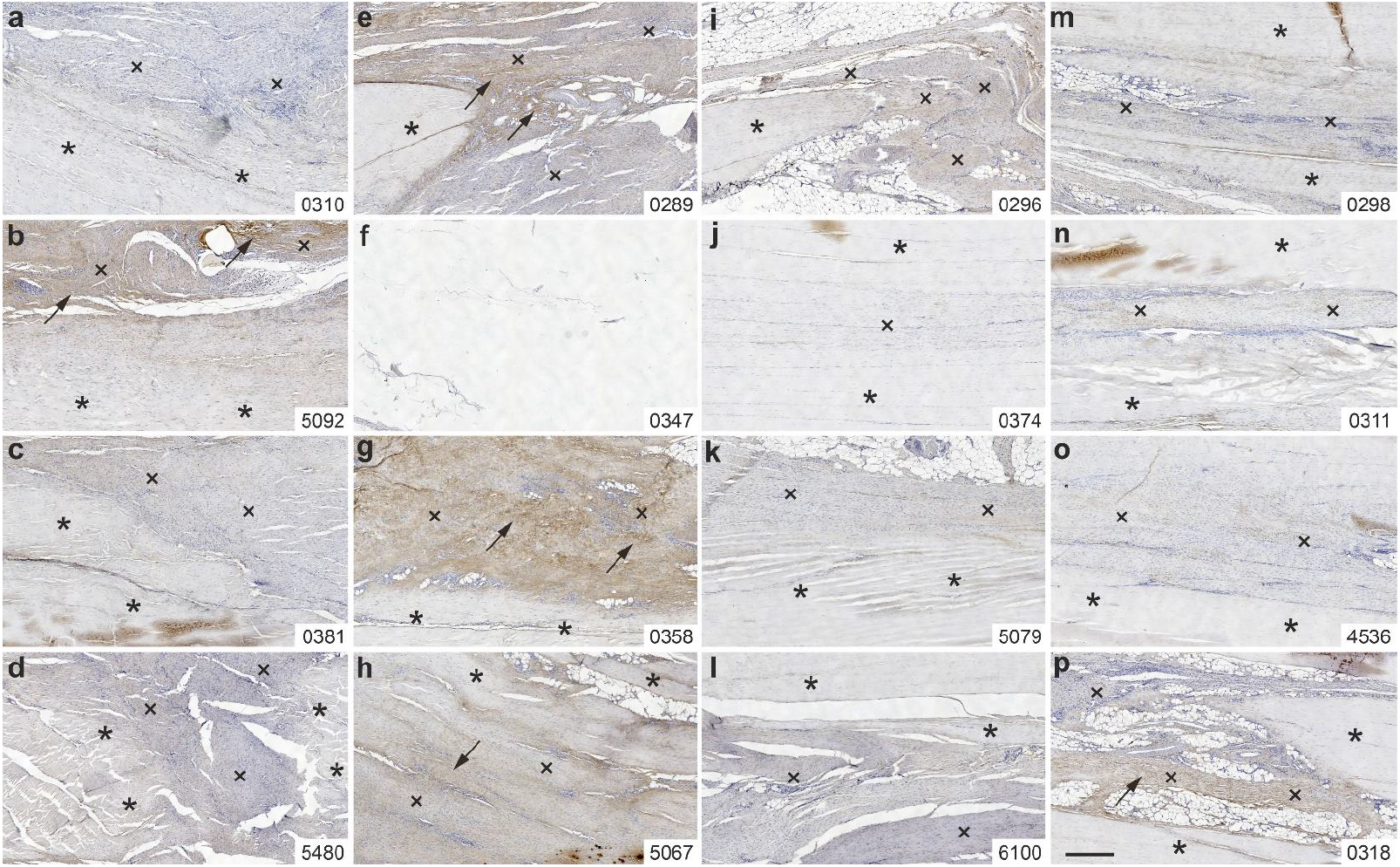
Representative photomicrographs of immunohistochemical detection of type III collagen in 5 μm thick sections of the right (surgery, treatment or sham treatment) common calcaneal tendon of the rabbits in Group 1 (UA-ADRCs / W4) (a-d), Group 3 (RLS / W4) (e-h), Group 2 (ADRCs / W12) (i-l) and Group 4 (RLS / W12) (m-p). The asterisks indicate original tendon tissue, the crosses point to newly formed connective tissue, and the arrows indicate immunolabeling for type III collagen in newly formed connective tissue. The numbers in the lower right corner of each panel identify the individual animals according to the study protocol. The scale bar in p represents 500 μm in all panels. Details are in the text.

One rabbit in Group 1 (UA-ADRCs / W4) (Fig. 10b) and three rabbits in Group 3 (RLS / W4) (Fig. 10e,g,h) showed immunolabeling for type III collagen in newly formed connective tissue (the section of rabbit 0347 (Group 3) processed for immunohistochemical detection of type III collagen did not show original tendon tissue and newly formed connective tissue and, thus, could not be evaluated). Furthermore, one rabbit in Group 4 (RLS / W12) showed discrete immunolabeling for type III collagen in newly formed connective tissue (Fig. 10p); the other animals in Group 4 as well as all animals in Group 2 (UA-ADRCs / W12) did not show immunolabeling for type III collagen.

### Immunolabeling of CD163

Figure 11 shows representative photomicrographs of immunohistochemical detection of CD163 in 5 μm thick sections of the right CCT of the rabbits in Group 1 (UA-ADRCs / W4) (Fig. 11a-d) and Group 3 (RLS / W4) (Fig. 11e-h).

**Fig. 11.**
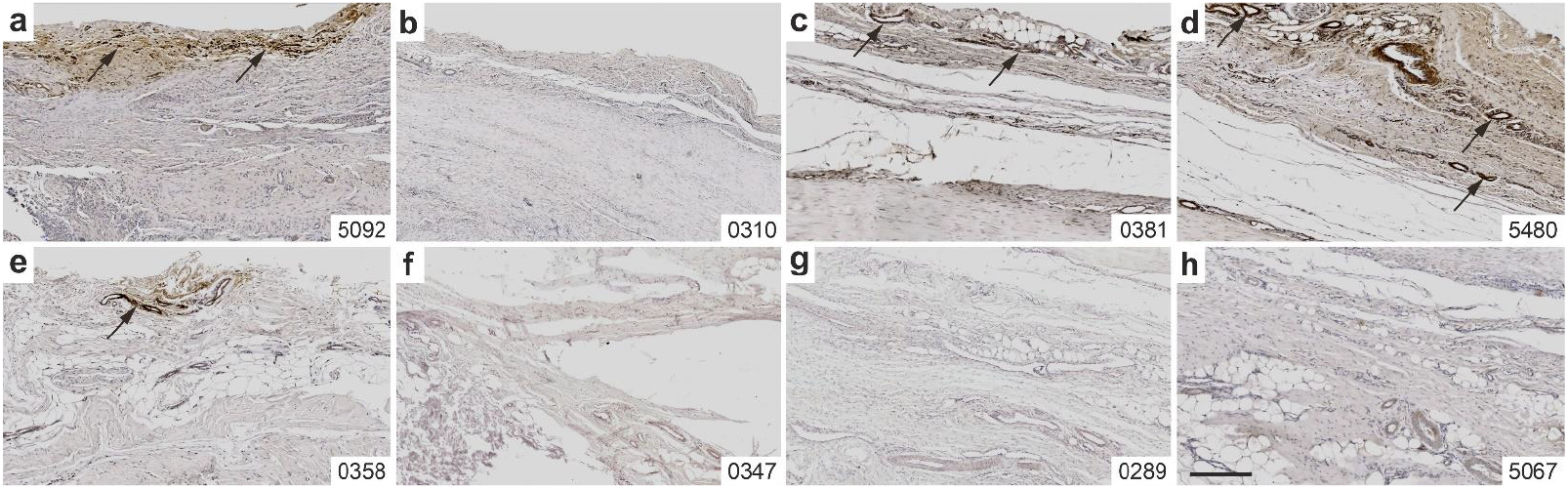
Representative photomicrographs of immunohistochemical detection of CD163 in 5 μm thick sections of the right (surgery, treatment or sham treatment) common calcaneal tendon of the rabbits in Group 1 (UA-ADRCs / W4) (a-d) and Group 3 (RLS / W4) (e-h). The arrows indicate detection of CD163 in paratenon tissue, mostly closely connected to vessels. The numbers in the lower right corner of each panel identify the individual animals according to the study protocol. The scale bar in h represents 300 μm in all panels. Details are in the text.

Immunolabeling for CD163 was found in the paratenon of the CCT of three animals in Group 1 (Fig. 11a,c,d) and, to a smaller extent, in the paratenon of the common calcaneal tendon of one animal in Group 3 (Fig. 11e). Immunolabeling for CD163 was most closely connected to vessels. No immunolabeling for CD163 was found in the paratenon of the right CCT of the animals in Group 2 (UA-ADRCs / W12) and Group 4 (RLS / W12) (photomicrographs not shown).

### Immunolabeling of aggrecan

Figure 12 shows overview photomicrographs of immunohistochemical detection of aggrecan in 5 μm thick sections of the right and left CCTs of all rabbits in Group 5 (UA-ADRCs / W12) (Fig. 12a,c) and Group 6 (RLS / W12) (Fig. 12b,d).

**Fig. 12.**
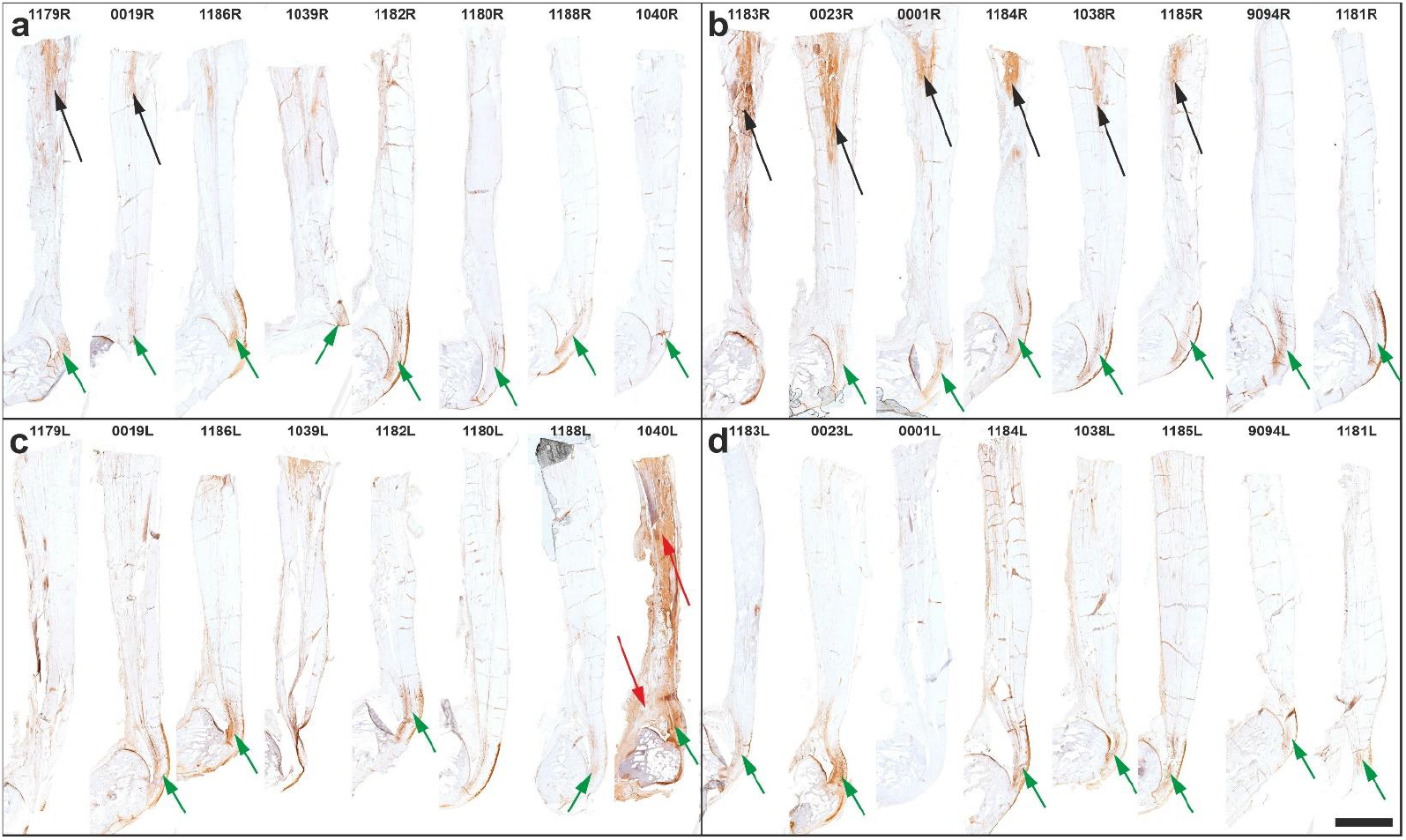
Representative photomicrographs of immunohistochemical detection of aggrecan in 5 μm thick sections of the right (surgery, treatment or sham treatment) (a,b) and the left (no surgery, no treatment or sham treatment) (c,d) common calcaneal tendon (CCT) of the rabbits in Group 5 (UA-ADRCs / W12) (a,c) and Group 6 (RLS / W12) (b,d). The green arrows indicate detection of aggrecan at the distal part of the CCT, the black arrows aggrecan at the site of surgery / treatment or sham treatment, and the red arrows a false positive immune reaction in the left CCT and paratenon tissue of rabbit 1040. The numbers above the photomicrographs identify the individual animals according to the study protocol. The scale bar in d represents 5 mm in all panels.

Sections of almost all rabbits showed the expected immunolabeling for aggrecan at the distal part of the CCT where the tendon is redirected by the posterosuperior corner of the calcaneus acting as a fulcrum, thus exposing the tendon to compressive loads (green arrows in Fig. 12). In addition, sections of six rabbits in Group 6 (RLS / W12) but only two rabbits in Group 5 (UA-ADRCs / W12) unexpectedly showed immunolabeling for aggrecan at the putative site of surgery and treatment or sham treatment (black arrows in Fig. 12).

### Combination of polarization microscopy and immunohistochemistry

Figure 13 shows representative polarization photomicrographs of sections stained with Safranin O/Fast Green (Fig. 13a-d, i-l) at the site of surgery and treatment as well as corresponding photomicrographs of immunohistochemical detection of aggrecan (Fig. 13e-h, m-p) of all rabbits in Group 5 (UA-ADRCs / W12); Figure 14 shows the corresponding photomicrographs of all rabbits in Group 6 (RLS / W12). In both figures the order of the animals (Panels a/e, b/f, c/g…l/p) is the same as in Figure 12 (from left to right).

**Fig. 13.**
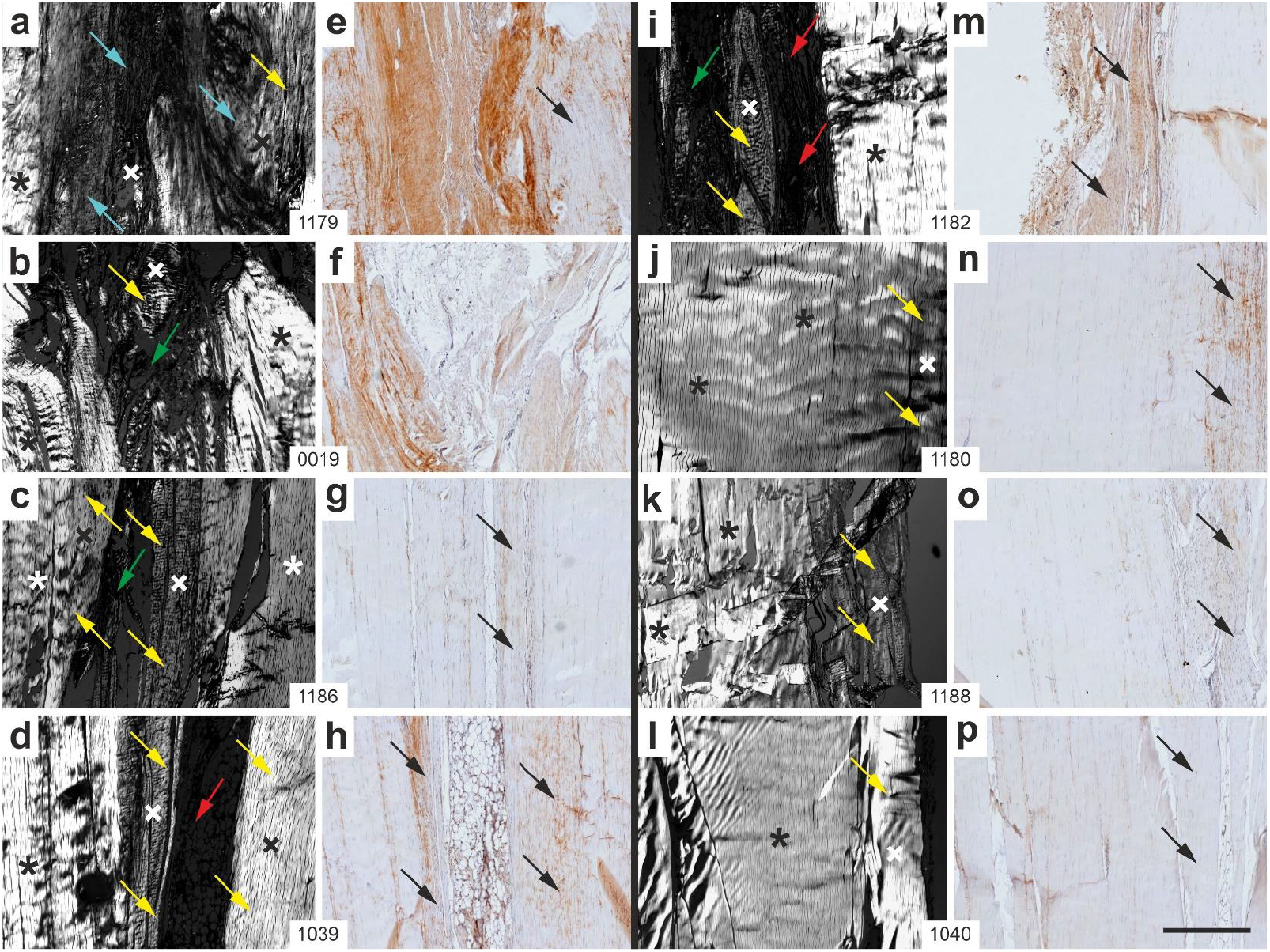
Representative polarization photomicrographs of 5 μm thick sections of the right common calcaneal tendon stained with Safranin O/Fast Green (a-d, i-l) at the site of surgery and treatment as well as corresponding photomicrographs of immunohistochemical detection of aggrecan (e-h, m-p) of all rabbits in Group 5 (UA-ADRCs / W12). The order of the animals (Panels a/e, b/f, c/g…l/p) is the same as in Figure 12a (from left to right). The asterisks indicate original tendon tissue, and the crosses point to newly formed connective tissue. The yellow arrows indicate newly formed connective tissue that showed the typical crimp pattern of tendons; the blue arrows point to newly formed connective tissue that did not show the typical crimp pattern of tendons; the green arrows indicate loose connective tissue and artifacts; the red arrows point to adipose tissue; and the black arrows indicate newly formed connective tissue in the photomicrographs of immunohistochemical detection of aggrecan that showed the typical crimp pattern of tendons in the polarization photomicropgraphs. The numbers in the white boxes that connect two corresponding panels each (a/e, b/f, c,g…l/p) identify the individual animals according to the study protocol. The scale bar in p represents 500 μm in all panels. Details are in the text.

**Fig. 14.**
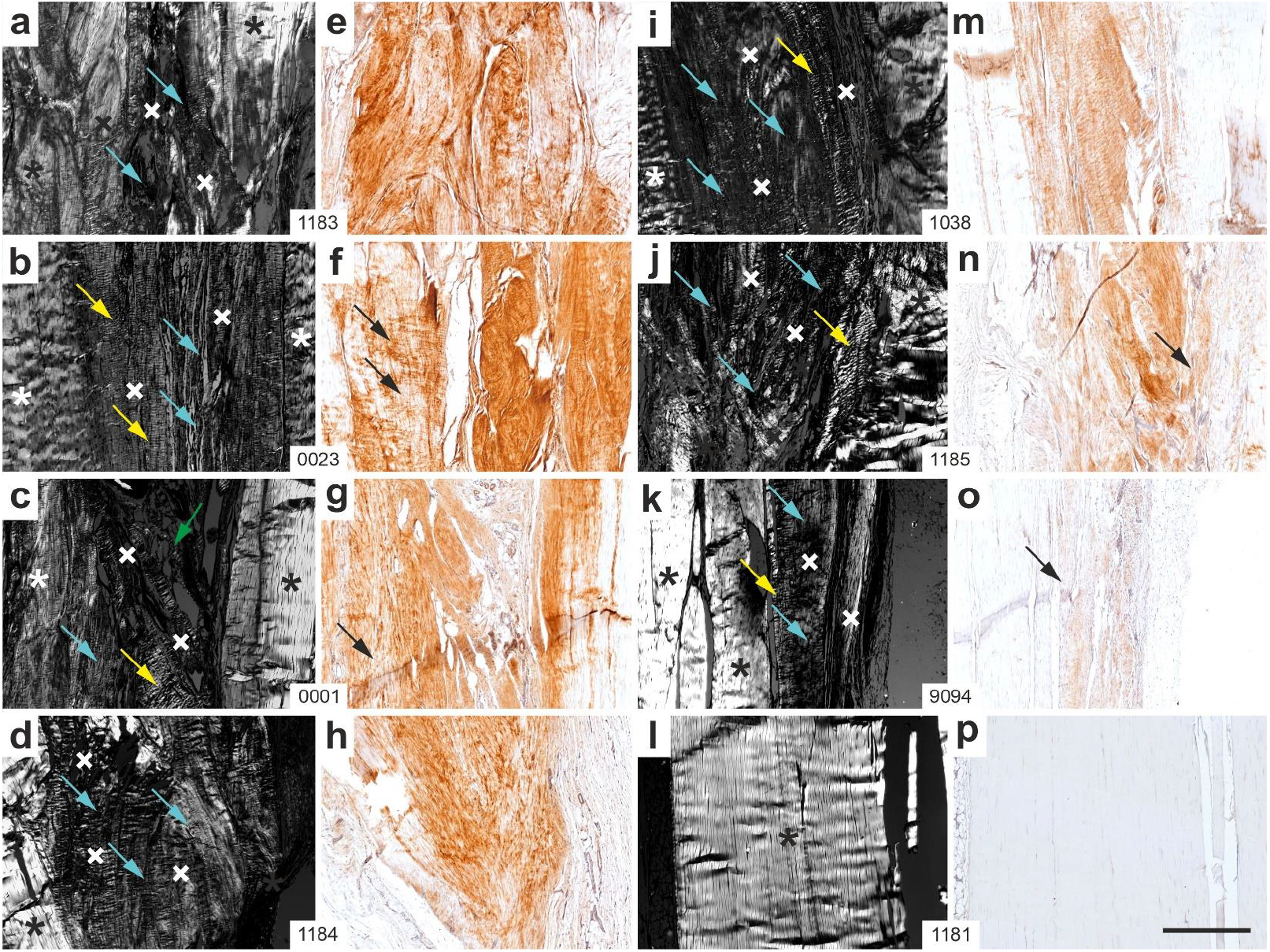
Representative polarization photomicrographs of 5 μm thick sections of the right common calcaneal tendon stained with Safranin O/Fast Green (a-d, i-l) at the site of surgery and sham treatment as well as corresponding photomicrographs of immunohistochemical detection of aggrecan (e-h, m-p) of all rabbits in Group 6 (RLS / W12). The order of the animals (Panels a/e, b/f, c/g…l/p) is the same as in Figure 12b (from left to right). The asterisks indicate original tendon tissue, and the crosses point to newly formed connective tissue. The yellow arrows indicate newly formed connective tissue that showed the typical crimp pattern of tendons; the blue arrows point to newly formed connective tissue that did not show the typical crimp pattern of tendons; the green arrows indicate loose connective tissue and artifacts; and the black arrows indicate newly formed connective tissue in the photomicrographs of immunohistochemical detection of aggrecan that showed the typical crimp pattern of tendons in the polarization photomicropgraphs. The numbers in the white boxes that connect two corresponding panels each (a/e, b/f, c,g…l/p) identify the individual animals according to the study protocol. The scale bar in p represents 500 μm in all panels. Details are in the text.

### Functional histology and functional immunohistochemistry

The results presented in this section represent functional histology and immunohistochemistry as the investigated distal parts of the CCT did not comprise the region of surgery and treatment or sham treatment.

Figure 15 shows anatomical details of the right distal CCT at the site of calcaneal insertion in a representative photomicrograph of a 5 μm thick section stained with Safranin O/Fast Green of a rabbit in Group 5 (UA-ADRCs / W12).

**Fig. 15.**
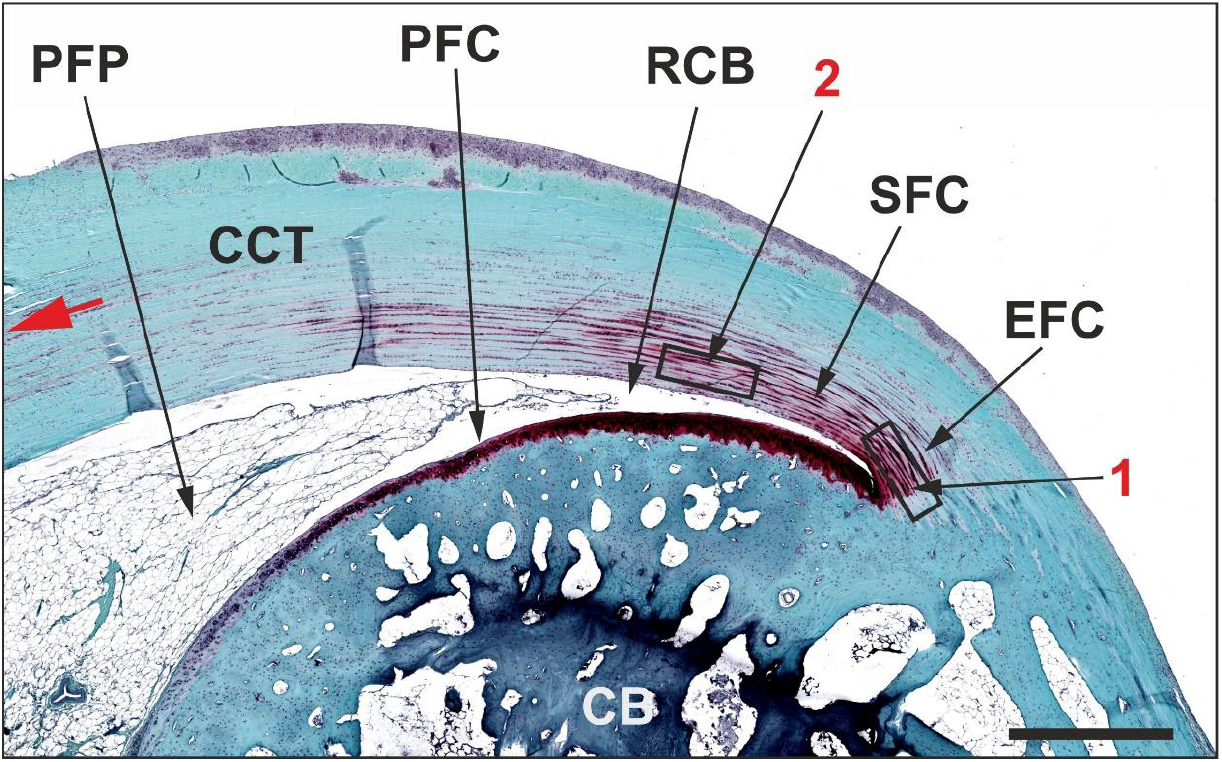
Anatomical details of the right common calcaneal tendon at the site of calcaneal insertion in a representative photomicrograph of a 5 μm thick section stained with Safranin O/Fast Green of a rabbit in Group 5 (UA-ADRCs / W12). Abbreviations: CB, calcaneal bone; CCT, common calcaneal tendon; ECF, enthesis fibrocartilage; PFC, periosteal fibrocartilage; PFP, precalcaneal fat pad; RCB, retro-calcaneal bursa; SFC, sesamoid fibrocartilage. The rectangles indicate the positions of the high-power photomicrographs of the CCT at the site of enthesis fibrocartilage (1) and sesamoid fibrocartilage (2) in Figs 17-19. The red arrow indicates the longitudinal axis of the CCT. The scale bar represents 1 mm.

Figure 16 shows representative low-magnification photomicrographs of 5 μm thick sections of the right, distal CCT at the site of calcaneal insertion stained with Safranin O / Fast Green (Fig. 16a-h, q-x) or processed for immunohistochemical detection of aggrecan (Fig. 16i-p,y-af) of all rabbits in Group 5 (UA-ADRCs / W12) (Fig. 16a-p) and Group 6 (RLS / W12) (Fig. 16q-af). The order of the animals (Panels a/i, b/j, c/k…x/af) is the same as in Figure 12a,b (from left to right).

**Fig. 16.**
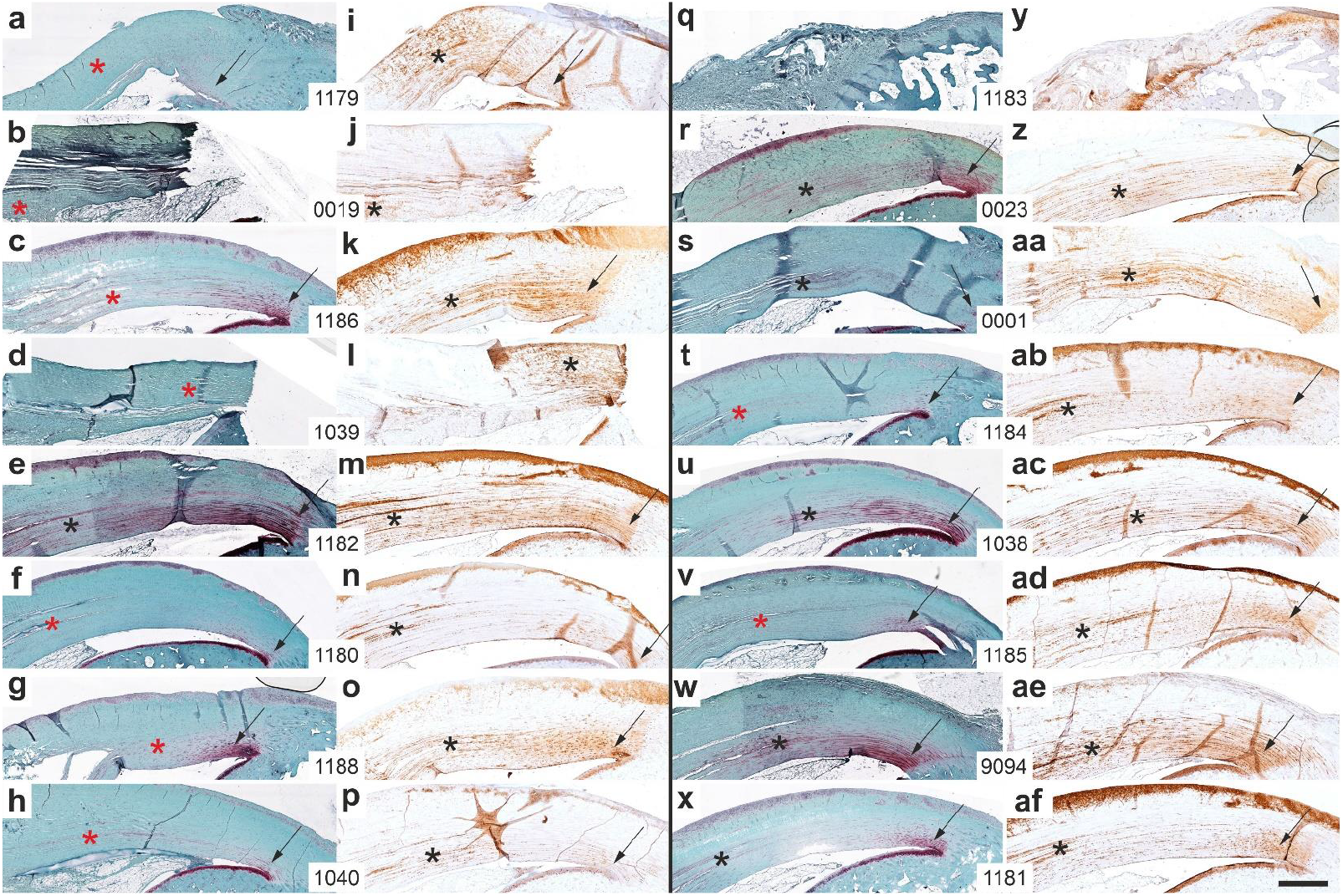
Representative low-magnification photomicrographs of 5 μm thick sections of the right (surgery, treatment or sham treatment), distal common calcaneal tendon (CCT) at the site of calcaneal insertion (similar to Fig. 15) stained with Safranin O / Fast Green (a-h, q-x) or processed for immunohistochemical detection of aggrecan (i-p,y-af) of all rabbits in Group 5 (UA-ADRCs / W12) (a-p) and Group 6 (RLS / W12) (q-af). The order of the animals (Panels a/i, b/j, c/k,…x/af) is the same as in Figure 12a,b (from left to right). The arrows indicate the transition from the CCT to the calcaneus. The asterisks indicate additional immunolabeling for aggrecan in the CCT at the site of the formation of sesamoid fibrocartilage as well as the corresponding regions in the Safranin O / Fast Green stained sections. The numbers in the white boxes that connect two corresponding panels each (a/i, b/j, c,k…x/af) identify the individual animals according to the study protocol. The scale bar in af represents 1 mm in all panels. Details are in the text.

In case of the rabbits 0019 (Fig. 16b,j) and 1039 (Fig. 16d,l) the CCT was torn from the calcaneus. Furthermore, in case of rabbit 1183 (Fig. 16q,y) the original CCT was not attached to the calcaneus, but the newly formed connective tissue was. Of note, although this newly formed connective tissue was firmly attached to the calcaneus, there was no expression of aggrecan which indicates that this newly formed tissue may have been functionally inactive.

The asterisks indicate additional immunolabeling for aggrecan in the CCT at the site of the formation of sesamoid fibrocartilage (Fig. 16i-p,z-af) as well as the corresponding regions in the Safranin O / Fast Green stained sections (Fig. 16a-h,r-x). One rabbit in Group 5 (rabbit 1182 in Group 5; Fig. 16e,m) and five rabbits in Group 6 (rabbits 0023 (Fig. 16r,z), 0001 (Fig. 16s,aa), 1038 (Fig. 16u,ac), 9094 (Fig. 16w,ae) and 1181 (Fig. 16x,af)) showed clear Safranin O staining at the location of the aggrecan signal (indicated by black asterisks in both corresponding panels each). In contrast, seven rabbits in Group 5 (all rabbits except rabbit 1182 (Fig. 16e,m)) but only two rabbits in Group 6 (rabbits 1184 (Fig. 16t,ab) and 1185 (Fig. 16v,ad) showed no or almost no Safranin O staining at the location of the aggrecan signal (indicated by red asterisks in the Safranin O stained panels in Fig. 16). This difference between Group 5 (UA-ADRCs / W12) and Group 6 (RLS / W12) was statistically significant (Fisher’s exact test; p = 0.041).

Figure 17 shows representative high-magnification photomicrographs of 5 μm thick sections of the right, distal CCT at the site of enthesis fibrocartilage (c.f. rectangle 1 in Fig. 15) stained with Safranin O / Fast Green (Fig. 17a-h, r-x) or processed for immunohistochemical detection of aggrecan (Fig. 17i-p,z-af) of all rabbits in Group 5 (UA-ADRCs / W12) (Fig. 17a-p) and Group 6 (RLS / W12) (Fig. 17r-af). The order of the animals (Panels a/i, b/j, c/k…x/af) is the same as in Figure 12a,b (from left to right).

**Fig. 17.**
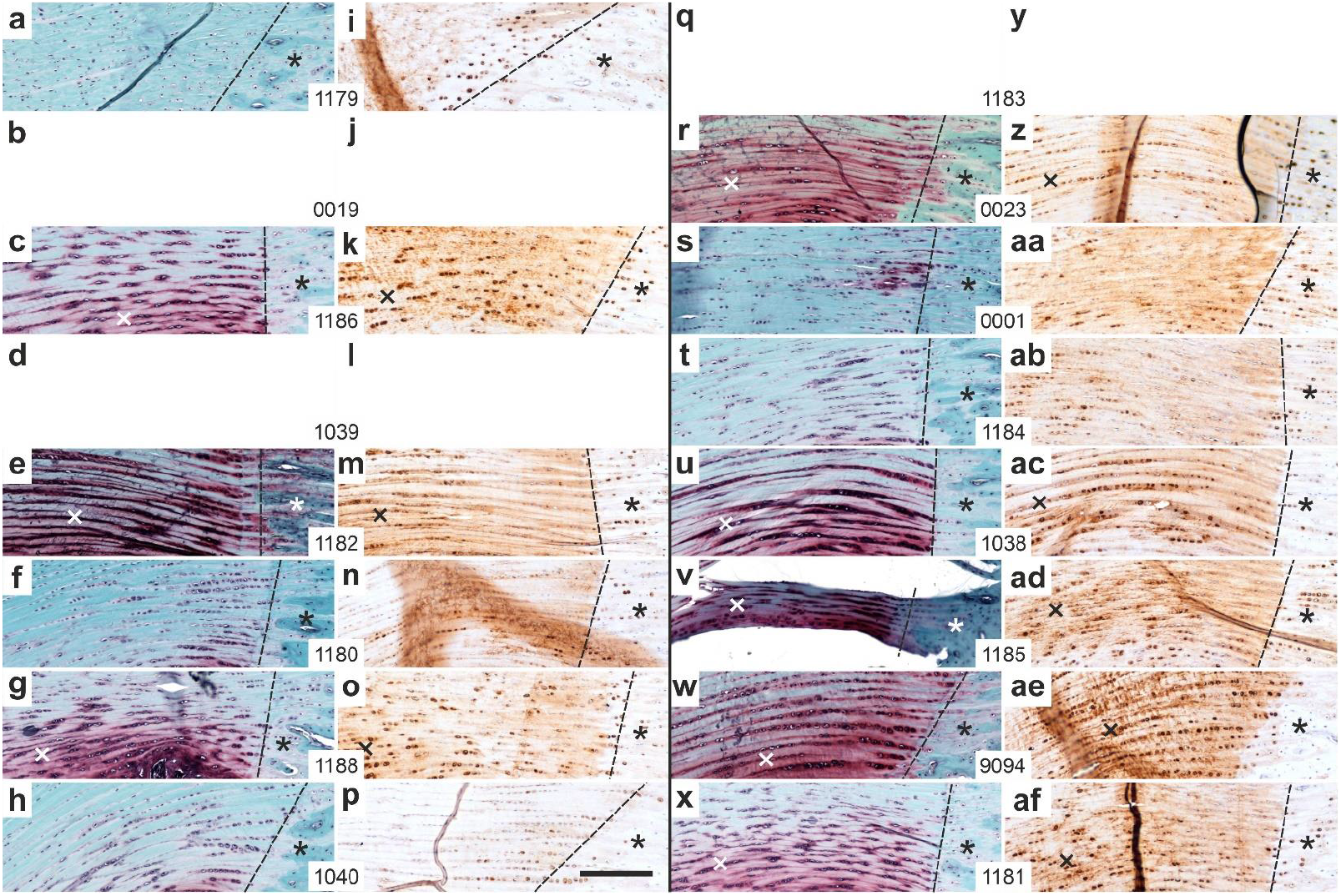
Representative high-magnification photomicrographs of 5 μm thick sections of the right (surgery, treatment or sham treatment), distal common calcaneal tendon (CCT) at the site of enthesis fibrocartilage (c.f. rectangle 1 in Fig. 15) stained with Safranin O / Fast Green (a-h, r-x) or processed for immunohistochemical detection of aggrecan (i-p,z-af) of all rabbits in Group 5 (UA-ADRCs / W12) (a-p) and Group 6 (RLS / W12) (r-af). The order of the animals (Panels a/i, b/j, c/k,…x/af) is the same as in Fig. 12a,b (from left to right). No photomicrographs of the right CCT of the rabbits 0019 (b,j) and 1039 (d,l) are shown as the CCT was torn from the calcaneus (c.f. Fig. 16b,j and d,l). Furthermore, no photomicrographs of the right CCT of rabbit 1183 (q,y) are shown as it was not the original CCT that was attached to the calcaneus, but rather newly formed connective tissue (c.f. Fig. 16q,y). The dashed lines indicate the transition from the CCT to the calcaneus (asterisks), and the crosses point to intense Safranin O staining of the enthesis fibrocartilage. The scale bar in p represents 200 μm in all panels.

It was not possible to clearly distinguish between the two groups.

Figure 18 shows representative high-magnification photomicrographs of 5 μm thick sections of the right, distal CCT at the site of sesamoid fibrocartilage (c.f. rectangle 2 in Fig. 15) stained with Safranin O / Fast Green (Fig. 18a-h, r-x) or processed for immunohistochemical detection of aggrecan (Fig. 18i-p,z-af) of all rabbits in Group 5 (UA-ADRCs / W12) (Fig. 18a-p) and Group 6 (RLS / W12) (Fig. 18r-af). The order of the animals (Panels a/i, b/j, c/k…x/af) is the same as in Figure 12a,b (from left to right).

**Fig. 18.**
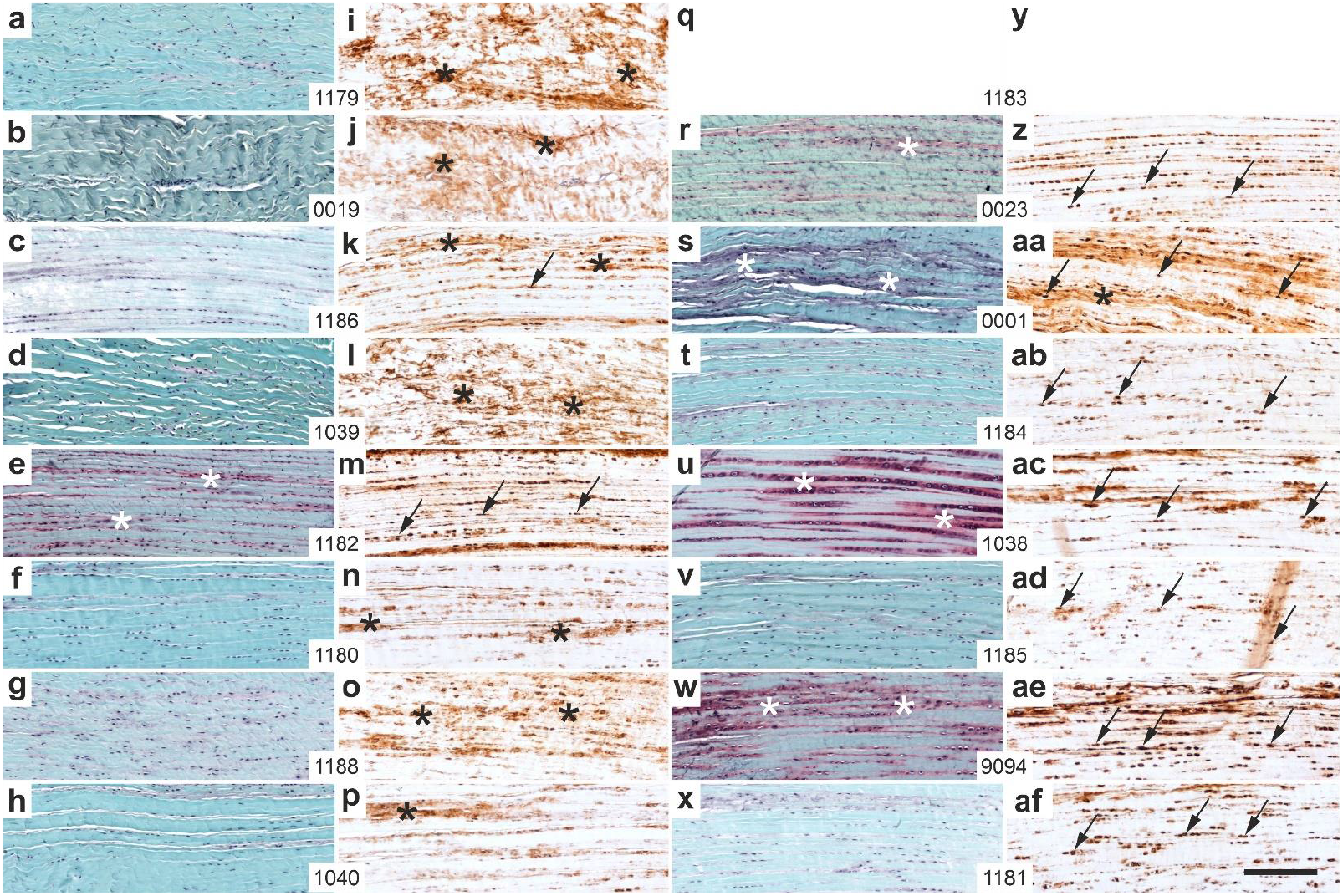
Representative high-magnification photomicrographs of 5 μm thick sections of the right (surgery, treatment or sham treatment), distal common calcaneal tendon (CCT) at the site of sesamoid fibrocartilage (c.f. rectangle 2 in Fig. 15) stained with Safranin O / Fast Green (a-h, r-x) or processed for immunohistochemical detection of aggrecan (i-p,z-af) of all rabbits in Group 5 (UA-ADRCs / W12) (a-p) and Group 6 (RLS / W12) (r-af). The order of the animals (Panels a/i, b/j, c/k,…x/af) is the same as in Figure 12a,b (from left to right). No photomicrographs of the right CCT of rabbit 1183 (q,y) are shown as it was not the original CCT that was attached to the calcaneus, but rather newly formed connective tissue (c.f. Fig. 16q,y). The arrows indicate intracellular immunolabeling for aggrecan, the black asterisks extracellular immunolabeling for aggrecan and the white asterisks intense Safranin O staining at the site of intracellular immunolabeling for aggrecan. The scale bar in af represents 200 μm in all panels.

Two rabbits in Group 5 (rabbis 1186 (Fig. 18k) and 1182 (Fig. 18m)), but all rabbits in Group 6, showed intracellular immunolabeling for aggrecan at the site of sesamoid fibrocartilage (arrows in Fig. 18). In contrast, except for rabbit 1182 all rabbits in Group 5, but only one rabbit in Group 6 (rabbit 0001; Fig. 18aa) showed intense, extracellular immunolabeling for aggrecan (black asterisks in Fig. 18). These differences between Group 5 (UA-ADRCs / W12) and Group 6 (RLS / W12) were statistically significant (Fisher’s exact test; p = 0.007 and 0.010).

In addition, the CCT of one rabbit in Group 5 (rabbit 1182; Fig. 18e) and four rabbits in Group 6 (rabbits 0012 (Fig. 18r), 0001 (Fig. 16s), 1038 (Fig. 18u) and 9094 (Fig. 18w)) showed intense Safranin O staining at the site of intracellular immunolabeling for aggrecan (white asterisks in Fig. 18).

Figure 19 shows representative high-magnification photomicrographs of 5 μm thick sections of the right and left, distal CCT at the site of enthesis fibrocartilage (c.f. rectangle 1 in Fig. 15) stained with Safranin O / Fast Green (Fig. 19a-h, q-x) or processed for immunohistochemical detection of aggrecan (Fig. 19i-p,y-af) of four rabbits each in Group 5 (UA-ADRCs / W12) (Fig. 19a-p) and Group 6 (RLS / W12) (Fig. 19q-af).

**Fig. 19.**
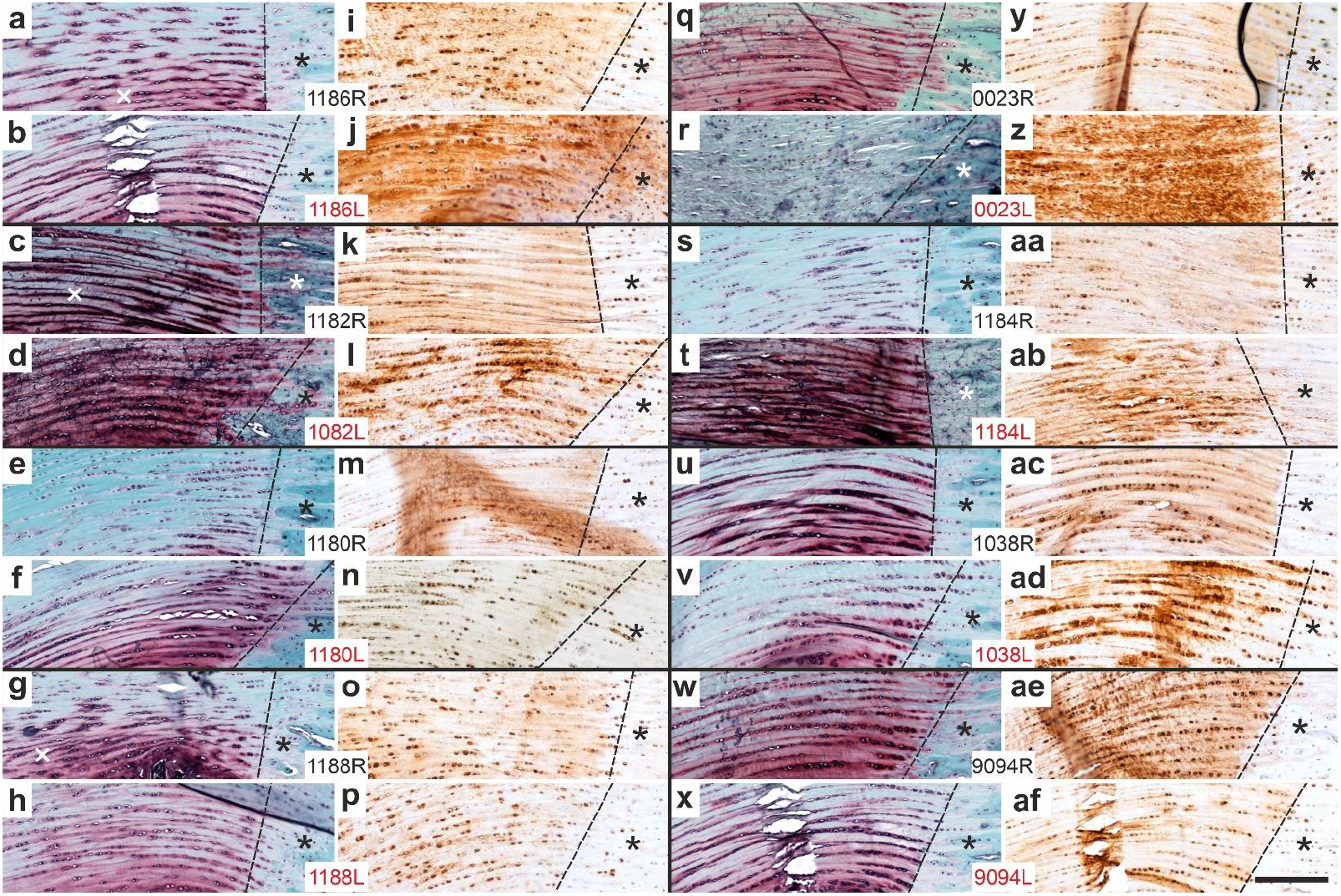
Representative high-magnification photomicrographs of 5 μm thick sections of the right (surgery, treatment or sham treatment) or left (no surgery, no treatment or sham treatment), distal common calcaneal tendon at the site of enthesis fibrocartilage (c.f. rectangle 1 in Fig. 13) stained with Safranin O / Fast Green (a-h, q-x) or processed for immunohistochemical detection of aggrecan (i-p,y-af) of four rabbits each in Group 5 (UA-ADRCs / W12) (a-p) and Group 6 (RLS / W12) (q-af). The dashed lines indicate the transition from the CCT to the calcaneus (asterisks). The scale bar in af represents 200 μm in all panels.

Furthermore, Figure 20 shows representative high-magnification photomicrographs of 5 μm thick sections of the right and left, distal CCT at the site of sesamoid fibrocartilage (c.f. rectangle 2 in Fig. 15) stained with Safranin O / Fast Green (Fig. 20-h, q-x) or processed for immunohistochemical detection of aggrecan (Fig. 20i-p,y-af) of four rabbits each in Group 5 (UA-ADRCs / W12) (Fig. 20a-p) and Group 6 (RLS / W12) (Fig. 20q-af).

**Fig. 20.**
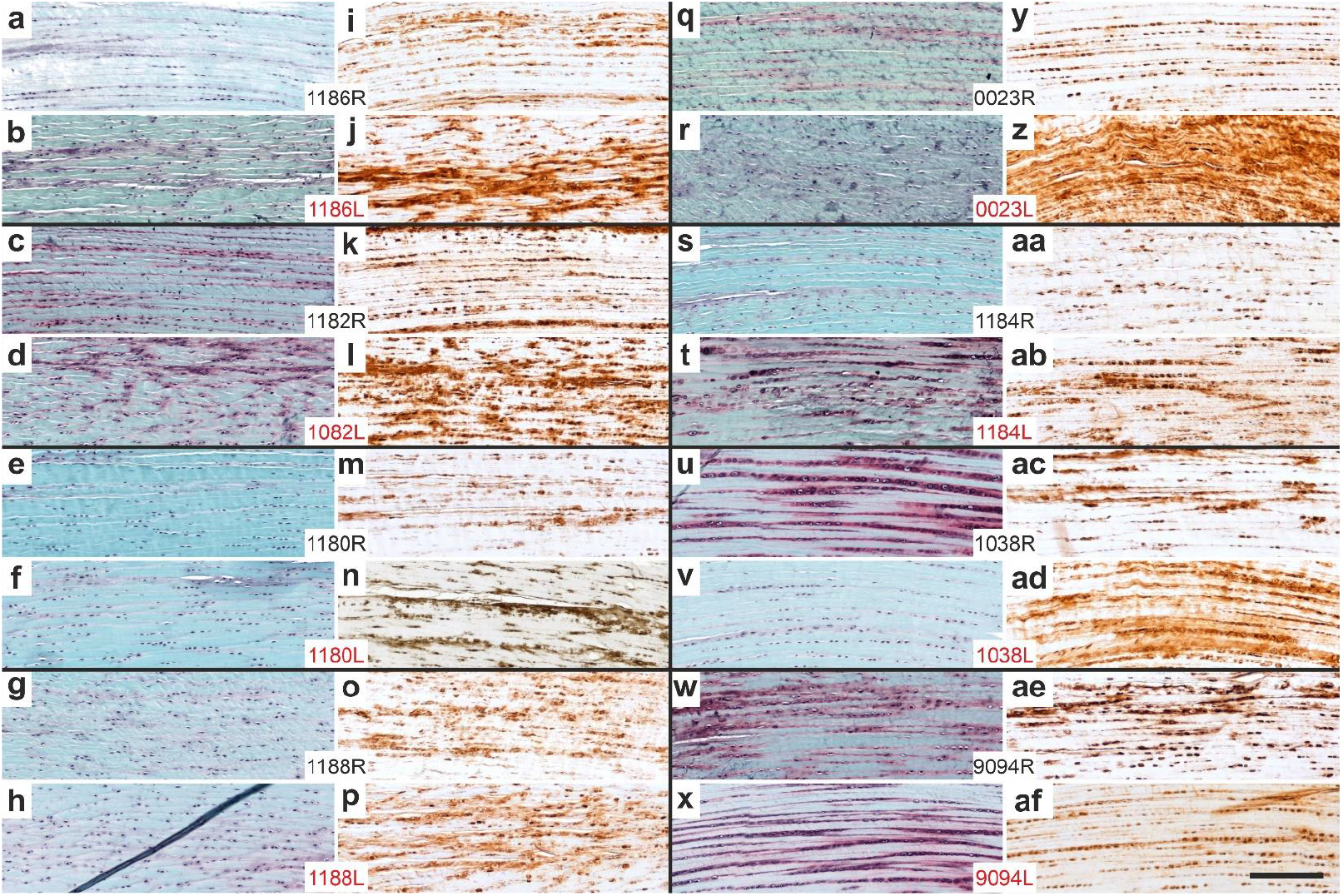
Representative high-magnification photomicrographs of 5 μm thick sections of the right (surgery, treatment or sham treatment) or left (no surgery, no treatment or sham treatment), distal common calcaneal tendon at the site of sesamoid fibrocartilage (c.f. rectangle 2 in Fig. 15) stained with Safranin O / Fast Green (a-h, q-x) or processed for immunohistochemical detection of aggrecan (i-p,y-af) of four rabbits each in Group 5 (UA-ADRCs / W12) (a-p) and Group 6 (RLS / W12) (q-af). The dashed lines indicate the transition from the CCT to the calcaneus (asterisks). The scale bar in af represents 200 μm in all panels.

Figures 19 and 20 show in both Groups 5 and 6 inter-individual differences in the Safranin O / Fast Green staining as well as the aggrecan immunolabeling patterns. These inter-individual differences might at least in part result from slight inter-individual differences in the anatomy and biomechanics of the CCT and the calcaneus (the more the CCT is “wrapped” around the calcaneus, the more lateral pressure is placed on the CCT). Of note, on average the intra-individual differences between the left and right CCTs of a given rabbit were more pronounced in Group 6 than in Group 5.

### Design-based stereologic analysis

Figure 21 shows the results of design-based stereologic analysis of the relative amount of cells, vessels, ECM and artifacts in the newly formed connective tissue of the sections stained with hematoxylin and eosine from the rabbits in Groups 1-4.

**Fig. 21.**
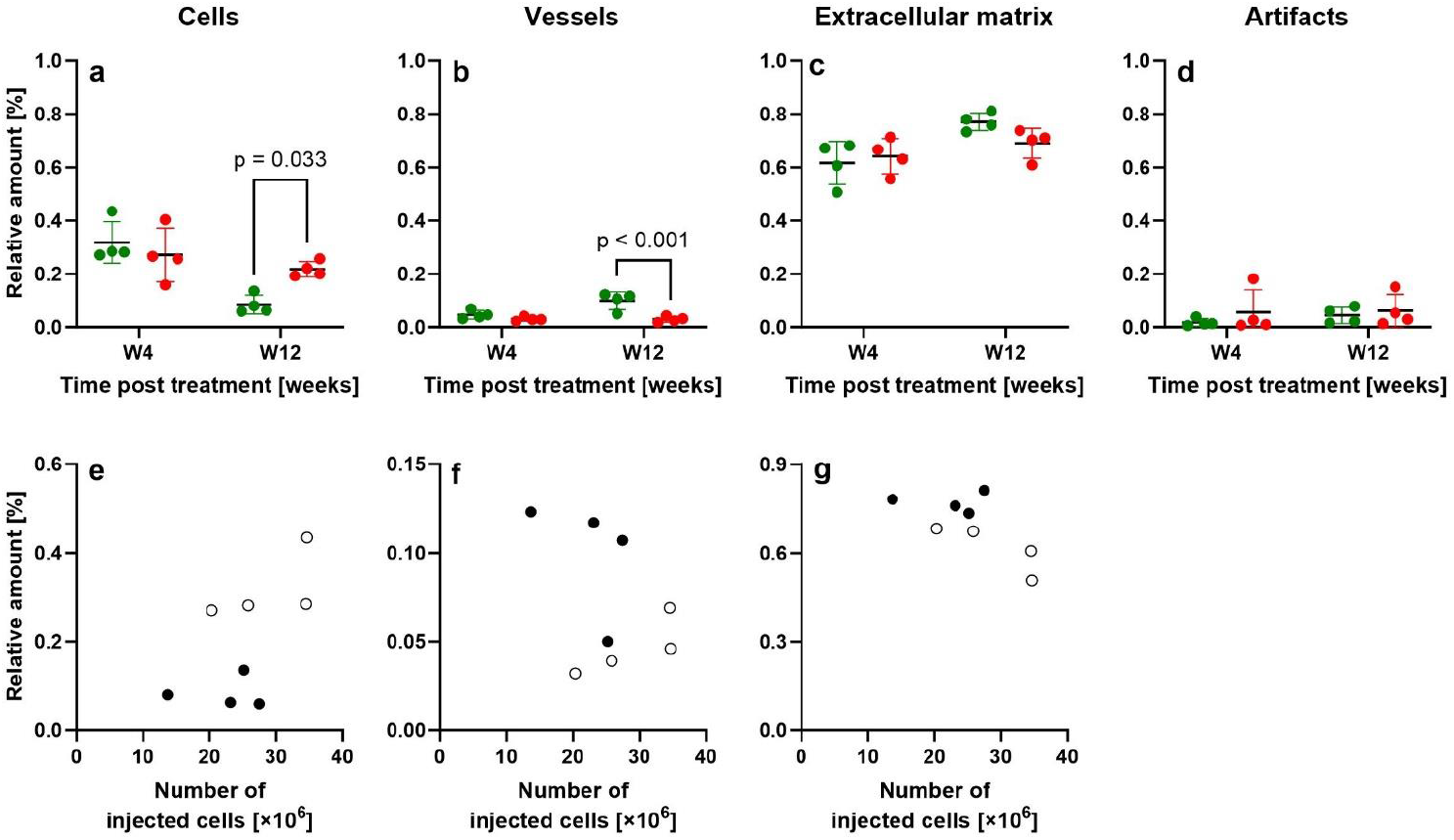
Scatter dot plots and mean ± standard deviation (horizontal lines) of the relative amount of cells (a), vessels (b), extracellular matrix (c), and artifacts (d) in the newly formed connective tissue of the right (surgery, treatment or sham treatment) common calcaneal tendon of the rabbits in Group 1 (UA-ADRCs / W4) (green dots at W4 in a-d; open dots in e-g), Group 3 (RLS / W4) (red dots at W4 in a-d), Group 2 (UA-ADRCs / W12) (green dots at W12 in a-d; closed dots in e-g) and Group 4 (RLS / W12) (red dots at W12 in a-d). The results of Bonferroni’s multiple comparison test with p < 0.05 are provided in a,b.

The results of two-way ANOVA (two treatments, two time points) are summarized in Table 3; the results of Bonferroni’s multiple comparison test for pairwise comparison are provided in Figure 21a-d.

**Table 3.**
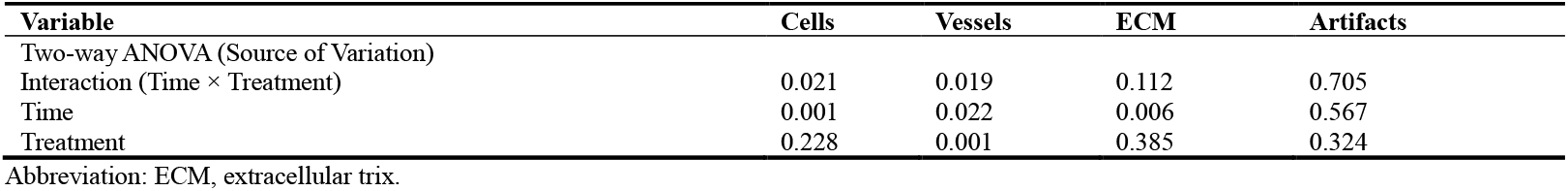
Results (p values) of the statistical analysis of the data shown in Figure 21 with two-way ANOVA.

There were no significant differences between the rabbits in Group 1 (UA-ADRCs / W4) and Group 3 (RLS / W4). However, the rabbits in Group 2 (UA-ADRCs / W12) showed a significantly lower relative amount of cells and a significantly higher relative amount of vessels in the newly formed connective tissue than the rabbits in Group 4 (RLS / W12) (p < 0.05). Of note, neither the rabbits in Group 1 nor the rabbits in Group 2 showed a significant correlation between the relative amount of cells, vessels and ECM and the initial cell dose (Fig. 21e-g).

### Bonar scores

Figure 22 shows the results of analyzing the newly formed connective tissue in the sections of the right CCT of the rabbits in Groups 1-4 using the Bonar score.

**Fig. 22.**
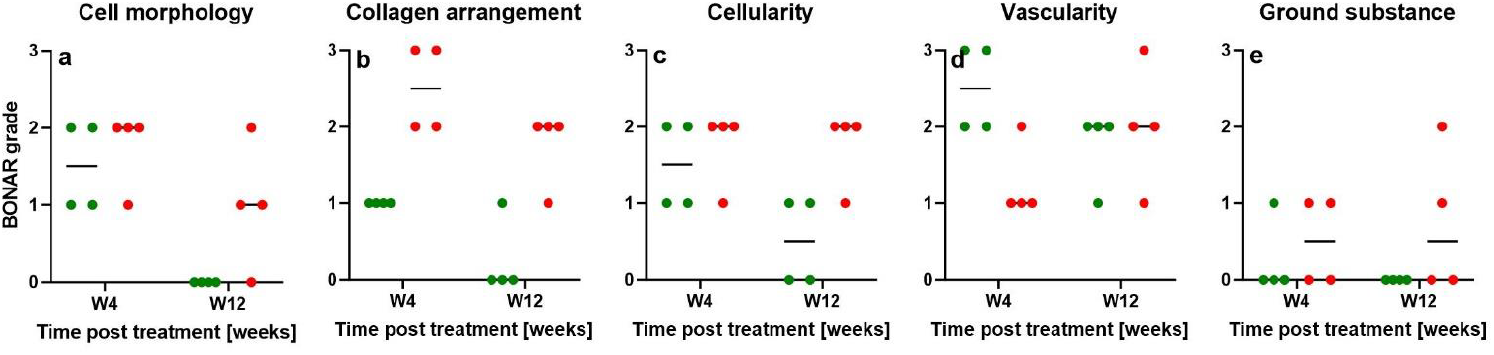
Scatter dot plots and median (horizontal lines) of the Bonar scores cell morphology (a), collagen arrangement (b), cellularity (c), vascularity (d) and ground substance (e) determined for the newly formed connective tissue in the sections of the right common calcaneal tendon of the rabbits in Group 1 (UA-ADRCs / W4) (green dots at W4 in a-e), Group 3 (RLS / W4) (red dots at W4 in a-e), Group 2 (UA-ADRCs / W12) (green dots at W12 in a-e) and Group 4 (RLS / W12) (red dots at W12 in a-e). The scores cell morphology, cellularity, vascularity and ground substance were determined on sections stained with hematoxylin and eosin, and the score collagen arrangement on sections stained with Picrosirius Red and evaluated using polarization microscopy.

Statistical analysis of these data with the Mann Whitney test did not show significant differences between the treatment with UA-ADRCs and the sham treatment (p>0.05 in all comparisons).

### Non-destructive biomechanical analysis

Figure 23a-e shows the results of the non-destructive biomechanical analysis of the left and right CCTs of the rabbits in Group 5 (UA-ADRCs / W12) and Group 6 (RLS / W12); Figure 23f shows the measurements of the cross-sectional area of these tendons.

**Fig. 23.**
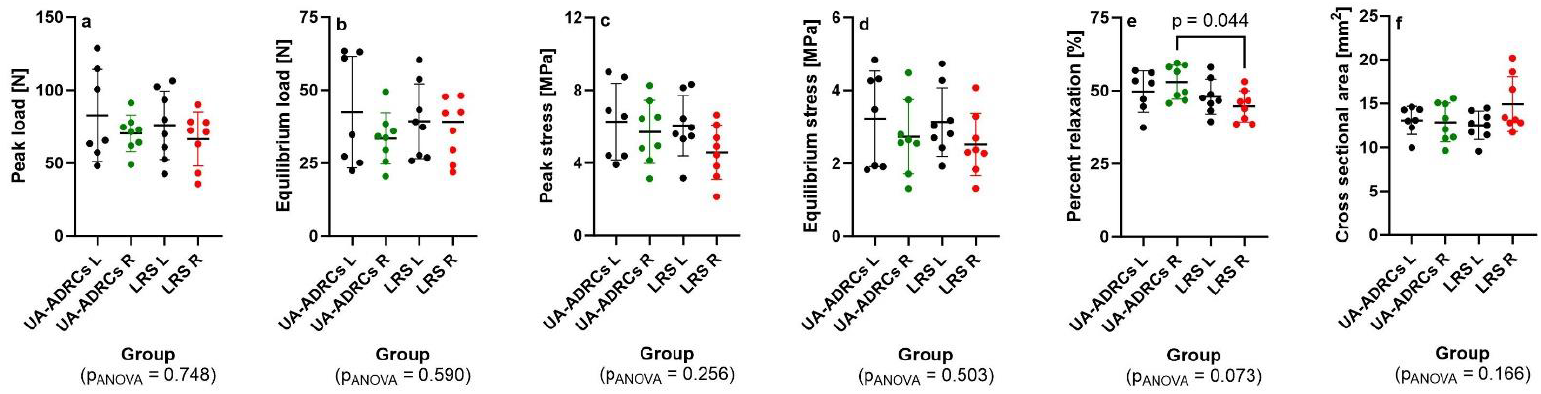
Scatter dot plots and mean ± standard deviation (horizontal lines) of the peak load (a), equilibrium load (b), peak stress (c), equilibrium stress (d), percent relaxation (e) and cross sectional area (f) of the left (intact / untreated) common calcaneal tendon of the rabbits in Group 5 (UA-ADRCs / W12) and Group 6 (RLS / W12) (black dots in a-f) as well as of the right common calcaneal tendon of the rabbits in Group 5 (green dots in a-f) and Group 6 (RLS / W12) (red dots in a-f). The results of one-way ANOVA are provided at the bottom of each panel, and the results of Bonferroni’s multiple comparison test with p < 0.05 in e.

Statistical analysis of these data with one-way ANOVA and Bonferroni’s multiple comparison test for pairwise comparison demonstrated a significantly smaller mean percent relaxation of the right (injured / sham treated) CCTs of the rabbits in Group 6 compared with the right (injured / treated) CCTs of the rabbits in Group 5 (Fig. 23e).

Figure 24 shows correlations between the individual results of the non-destructive biomechanical analysis and the cross-sectional area of the CCT of all rabbits in Group 5 (UA-ADRCs / W12) and Group 6 (RLS / W12).

**Fig. 24.**
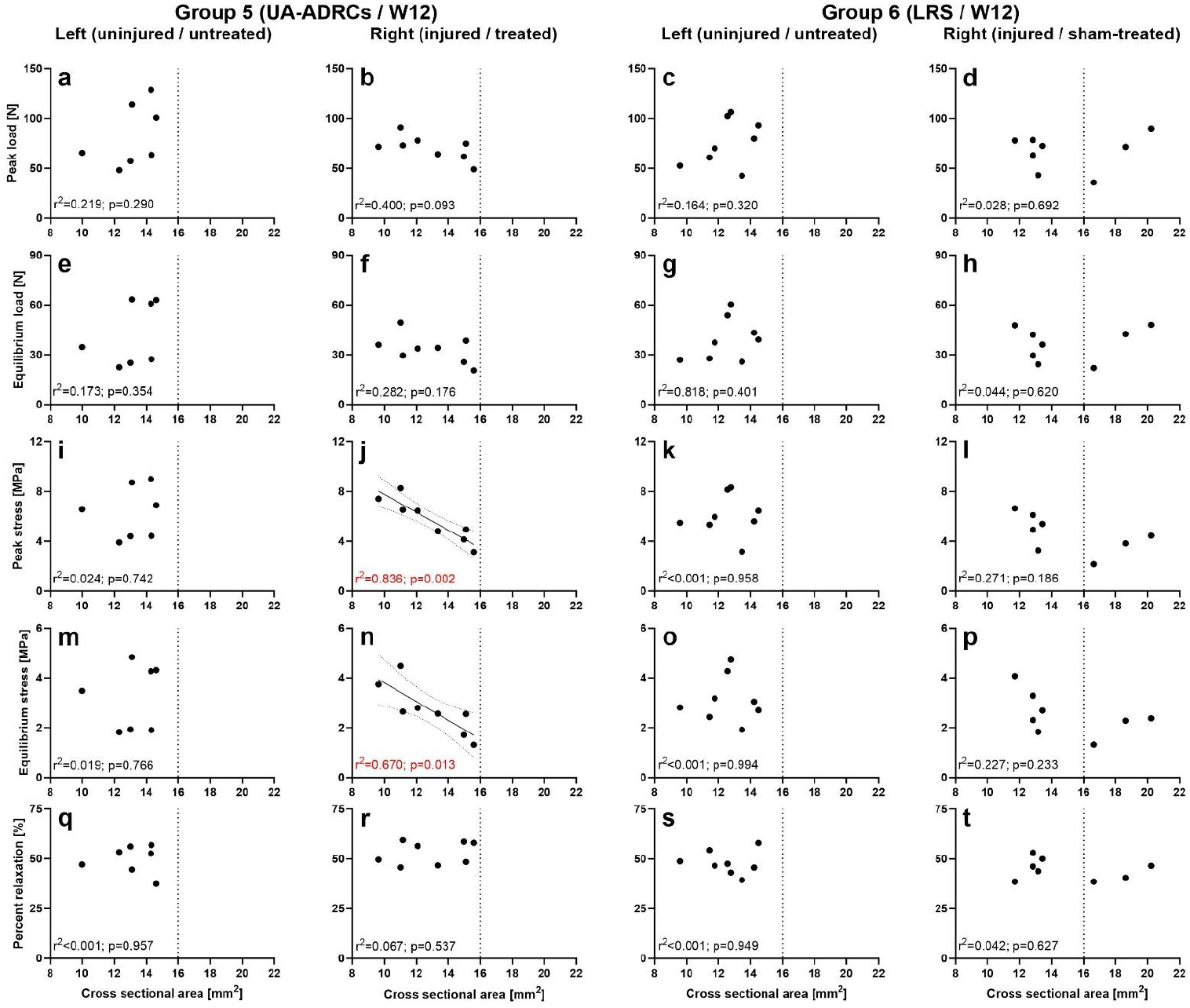
Scatter dot plots and results of linear regression analysis of the peak load (a-d), equilibrium load (e-h), peak stress (i-l), equilibrium stress (m-p) and percent relaxation (q-t) as a function of the cross sectional area of the left (uninjured / untreated) common calcaneal tendon (CCT) of the rabbits in Group 5 (UA-ADRCs / W12) (a,e,i,m,q) and Group 6 (RLS / W12) (c,g,k,o,s) as well as of the right common calcaneal tendon of the rabbits in Group 5 (UA-ADRCs / W12) (b,f,j,n,r) and Group 6 (RLS / W12) (d,h,l,p,t). The results of linear regression analysis (coefficient of determination (r2) and p-value) are provided in each panel. In case of p < 0.05 the best-fit lines (solid lines in j,n) together with their 95% confidence bands (dotted curves in j,n) are indicated. The vertical dotted lines in all panels represent a cross sectional area of 16 mm2. Three rabbits in Group 6 had a right CCT with cross-sectional area >16 mm2 (d,h,l,p,t); all other CCTs had a cross-sectional area < 16 mm2.

Significant, negative correlations between the peak stress and the cross-sectional area of the CCT as well as between the equilibrium stress and the cross-sectional area of the CCT were found after treatment with UA-ADRCs (Fig. 24j,n) but not after sham treatment (Fig. 24l,p). No other significant correlations between the results of the non-destructive biomechanical analysis and the cross-sectional area of the CCT were found. Of note, there were no significant correlations between the results of the biomechanical analysis and the initial cell dose (Fig. 25a-f).

**Fig. 25.**
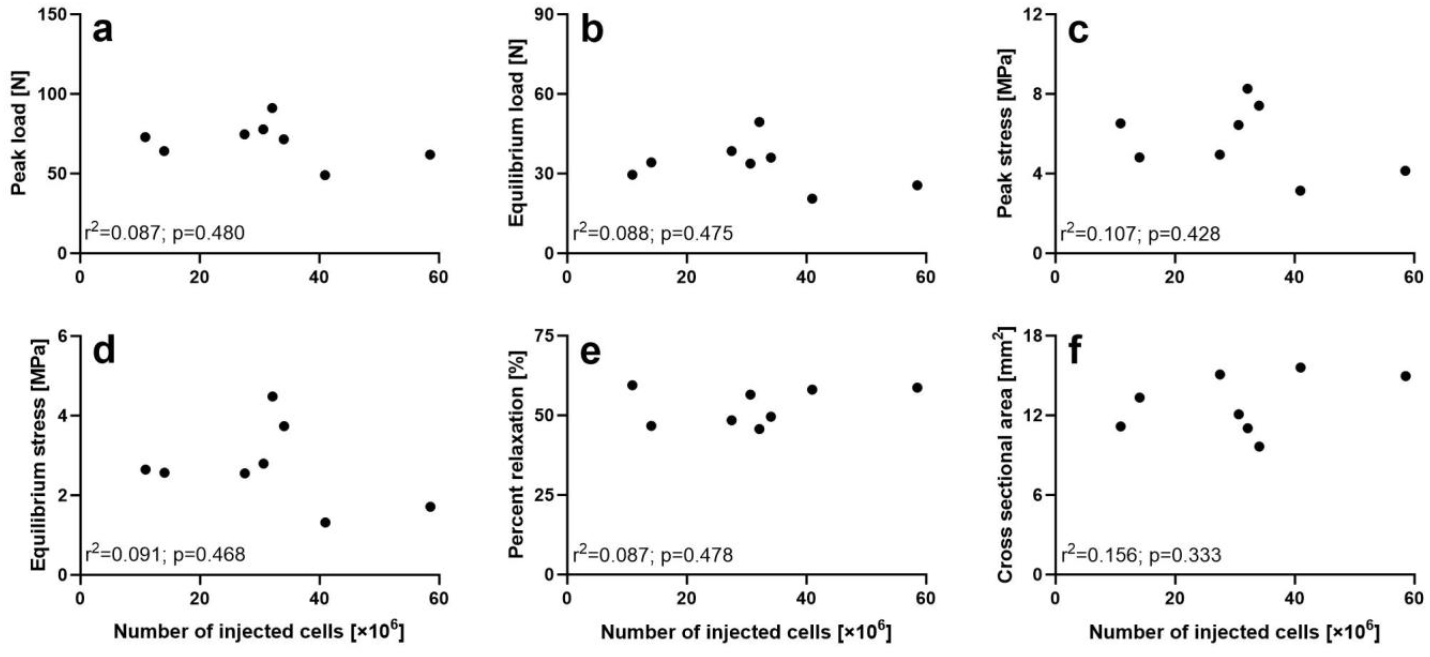
Scatter dot plots and results of linear regression analysis of the peak load (a), equilibrium load (b), peak stress (c), equilibrium stress (d), percent relaxation (e) and the cross sectional area of the right common calcaneal tendon of the rabbits in Group 5 (UA-ADRCs / W12) as a function of the number of injected cells. The results of linear regression analysis (coefficient of determination (r2) and p-value) are provided in each panel.

## DISCUSSION

In principle, UA-ADRCs cannot be labeled and, thus, not detected in the host tissue [20,21]. Furthermore, molecular and cellular mechanisms of actions of UA-ADRCs cannot be investigated in vitro as the composition of UA-ADRCs immediately changes right after plating for expansion and culture [22]. The latter results in adipose derived stem cells (ADSCs) [23] that have been extensively investigated in the literature (e.g. [24-26]), are capable of differentiating into somatic cells of the three germ layers (e.g. [3,21]) and can be detected in the host tissue after labeling with e.g, green fluorescence proteine (shown for autologous cultured ADSCs in e.g. [20] and for allogeneic cultured ADSCs in e.g. [27]). However, these stem cells only represent a small relative number of the cells contained in UA-ADRCs [5].

Therefore all conclusions regarding the effects of UA-ADRCs in the host tissue must be drawn from observed alterations of the structure and function of the host tissue after treatment with UA-ADRCs in comparison with sham treatment or an alternative treatment.

To our knowledge, this is the first study which provides comprehensive insights into the histological, immunohistochemical and biomechanical properties of a tendon after experimental induction of a partial-thickness tear followed by treatment with UA-ADRCs in vivo. The motivation for using the different staining, immunohistochemical labeling and microscopic techniques in this investigation was as follows: (i) Tendons are composed of longitudinally aligned type I collagen fibrils arranged in bundles with an undulating pattern. This pattern, which is called crimp, is established during fetal development [28]. The crimp structure that can be visualized using polarization microscopy plays a pivotal role in the mechanical behavior of tendons, acting as a shock absorber during loading [29]. (ii) Type I procollagen is a triple-stranded, rope-like molecule that is processed by enzymes outside the cell, followed by self-arrangement of the processed molecules into long, thin type I collagen fibrils that cross-link to one another in the extracellular space [30,31]. (iii) Unorganized type III collagen is present in scar tissue and represents higher hardness but lower strength than organized type I collagen [32,33]. (iv) CD163 is usually considered a marker of M2 macrophages [34], which are mainly involved in anti-inflammatory responses [35]. Tears of the rotator cuff are associated with synovial inflammation and increased expression of the pro-inflammatory markers interleukin-1β (IL-1β) and tumor necrosis factor-α (TNF-α) [36,37] (for the role of low-grade inflammation in the pathogenesis of tendinopathy see also [38]). Furthermore, the exposure of cultured, adipose-derived stem cells (ADSCs) with IL-1β and TNF-α results in decreased expression of the tenogenic transcription factor scleraxis [39]. The latter plays a pivotal role in promoting tenocyte proliferation and the synthesis of extracellular matrix (ECM) during embryonic tendon development, and is also involved in promoting the initial expansion of newly committed tenocytes and the production of ECM proteins in adult tendons [40]. This is in line with the finding that exposure of postnatal tendon cells with IL-1β resulted in reduced anabolic activity (leading to abnormal ECM deposition and organization) and increased catabolic activity (leading to proinflammatory cues and ECM degradation) [41]. In summary, M2 macrophages in injured tendon tissue may significantly contribute to tendon regeneration. (v) Aggrecan is a marker of tissues with a fibrocartilaginous phenotype and thus, intermittent compressive load acting on the tendinous tissue [42]. The presence of aggrecan in a tendon specifically increases its capacity to imbibe water and, thus, to withstand compression [43]. (vi) Safranin O binds to proteoglycans [44] and stains them bright red; these proteoglycans serve as water storage [45] (similar to aggrecan). Fast Green stains collagen.

Our results indicate seamless integration of newly formed connective tissue with characteristics of tendon tissue into the existing tendon tissue, with biomechanical properties pointing to homogeneous tissue composition (indicated by a significant, negative correlation between the peak stress and the cross-sectional area of the CCT as well as between the equilibrium stress and the cross-sectional area of the CCT after treatment with UA-ADRCs). In contrast, sham treatment resulted in the formation of tissue with characteristics of scar tissue, without seamless integration of the newly formed connective tissue into the existing tendon tissue but the formation of blood vessels and adipose tissue at the junction, and no correlation between the peak stress and the cross-sectional area of the CCT as well as between the equilibrium stress and the cross-sectional area of the CCT. Furthermore, treatment with UA-ADRCs resulted in a significantly higher mean elasticity of the CCT (indicated by the percent relaxation) than sham treatment.

Collectively, four main findings emerge from the complex data shown in Figs 16-20: (i) Aggrecan was primarily found in the newly formed connective tissue that did not show the typical crimp pattern of tendons. In line with the well-known molecular parameters indicating adaptation to mechanical stress in fibrous connective tissue (reviewed in [15,43]) the irregular arrangement of the collagen fibers in the newly formed connective tissue that did not show the typical crimp pattern of tendons might have put the tendon cells under pressure and, thus, initiate the expression of aggrecan.

(ii) Intracellular aggrecan cannot serve as a water reservoir in the ECM. In this case the proteoglycans (detected by Safranin O staining) may have taken over this function in the CCT, with more rabbits in Group 6 than in Group 5 having shown this phenomenon. (iii) Overall, the rabbits in Group 6 lagged behind in aggrecan expression, which could indicate that the right (injured / sham treated) hindlimb of the rabbits in Group 6 was functionally impaired for longer than the left (uninjured / untreated) hindlimb of these rabbits. (iv) The right (injured / treated) CCT of the rabbits in Group 5 (as far as can be judged from Safranin O / Fast Green staining and immunolabeling for aggrecan) had largely returned to their normal function after 12 weeks post-treatment, which was not the case with the right (injured / sham treated) CCT of the rabbits in Group 6.

Of particular importance was the substantial reduction in the relative amount of cells in the newly formed connective tissue between W4 and W12 post-treatment after treatment with UA-ADRCs (−73%) but not after sham treatment (−20%) (Fig. 21a). In this regard a recent study on the development of mouse tendon tissue indicated that the time interval between postnatal day 4 (P4) and P12 is the critical stage for postnatal tendon maturation [46]. Specifically, this study demonstrated that the gene expression of scleraxis, mohawk, type IA1 collagen and tenomodulin peaked at P4, followed by a substantial reduction in the relative amount of cells in the tendon tissue between P14 and P28 [46]. In a study on human fetal posterior tibial tendons samples obtained from 22-28 week old fetuses showed a higher relative amount of cells in the tendon tissue than samples obtained from 32-38 week old fetuses [28]. Taking this into account, treatment of a partial-thickness tear of a tendon with UA-ADRCs might lead to tissue regeneration that shares certain characteristics with late fetal / early postnatal tendon maturation.

Another key finding of the present study was the formation of adipose tissue in the defective area of the tendon after sham treatment, but not after treatment with UA-ADRCs (Fig. 6). We had already observed a similar phenomenon in a first-in-human case report of guided bone regeneration / maxillary sinus augmentation in oral surgery prior to implant placement, with a higher relative amount of newly formed bone at six weeks (W6) post-treatment with UA-ADRCs, fraction 2 of plasma rich in growth factors (PRGF-2) and an osteoinductive scaffold (OIS) (UA-ADRCs/PRGF-2/OIS) than at W34 post-treatment with PRGF-2/OIS alone (W6: 20.2% vs 12.4%; W34: 24.2% vs 16.8%), and a higher relative amount of adipose tissue after treatment with PRGF-2/OIS than after treatment with UA-ADRCs/PRGF-2/OIS (W6: 0.2% vs 0%; W34: 15% vs 2.6%) [47]. In this regard it is crucial to understand that ADRCs are neither ‘fat stem cells’ nor could they exclusively be isolated from adipose tissue [20,21]. Rather, ADRCs contain the same adult stem cells that are ubiquitously present in the walls of small blood vessels, capable to differentiate into somatic cells of the three germ layers (c.f. [3,21]).

The results of the present study are also in line with our observations on the biopsy of a supraspinatus tendon of a patient with traumatic rotator cuff injury, taken ten weeks after local injection of UA-ADRCs [11]. Similar to the present study we observed newly formed, well organized, firm connective tissue with discernible crimp arrangement and immunohistochemical detection of type I procollagen, with no formation of adipocytes in the biopsy [11]. This was accompanied by immunohistochemical detection of abundant, intracellular and extracellular tenomodulin at the putative injection site, as well as immunohistochemical detection of the proliferation marker Ki-67 in cells with the characteristic morphology of tenocytes [11].

To our knowledge treatment of experimentally induced tendon injuries with UA-ADRCs in vivo has only been addressed in two studies to date [48,49]. In these studies the supraspinatus tendon of rabbits was severed from the greater tuberosity, followed by suturing the severed supraspinatus tendon to the tuberosity through the bone and application of UA-ADRCs in fibrin glue or fibrin glue alone as control treatment [48,49]. The authors reported higher levels of type I collagen and a higher type I / type III collagen ratio after treatment with UA-ADRCs than after sham treatment, as well as improved biomechanical properties (significantly higher mean maximum load, mean maximum stiffness and mean maximum strength after treatment with UA-ADRCs than after control treatment) [48,49]. Although not directly comparable, the results reported in [48,49] are consistent with the results of the present study.

Another study evidenced significantly higher gene expression of type IA1 collagen and vascular endothelial growth factor (VEGF) in tendon tissue after treatment of a collagenase-induced sheep Achilles tendinopathy model with autologous, microfragmented fat compared to sham treatment, but significantly lower gene expression of scleraxis, TGF-ß1, type IIIA1 collagen, matrix metalloproteinase 1 (MMP1), MMP9 and interleukin 1 [50]. Biomechanical analysis showed no significant difference between treatment with microfragmented fat and sham treatment [50]. In this regard, autologous, microfragmented fat cannot be compared with UA-ADRCs (outlined in detail in [5]). Specifically, in contrast to UA-ADRCs, microfragmented fat contains adipocytes and connective tissue (c.f. Fig. 3 in [51]), and many cells of the stromal vascular fraction may be lost during filtration steps because they are not released from connective tissue fragments (c.f. [3,5]).

A clinical study found no improvement of anterior cruciate ligament reconstruction by injection of UA-ADRCs into a bone-tendon-bone (BTB) graft derived from the patella and related tendons compared without injection of UA-ADRCs [46]. However, the description of the infiltration procedure of the BTB graft with UA-ADRCs in that study [52] does not exclude the cells were injected mostly into the tendon part of the BTB graft, which may have limited the success of their approach.

Treatment of experimentally induced tendon injuries with autologous, cultured adipose derived stem cells (ADSCs) in vivo was addressed in several studies (Table 4; [7,53-68]).

**Table 4.** Details of studies on treatment of experimentally induced tendon injuries with autologous, cultured, adipose derived stem cells in vivo.

| R | Y | Sp | T | I | P | S | AT | C | H | PM | IHC | B |
| --- | --- | --- | --- | --- | --- | --- | --- | --- | --- | --- | --- | --- |
| [53] | 2011 | Rabbit | CCT | Incision | + | - | PRP | - | + | - | - | - |
| [54] | 2012 | Rabbit | CCT | Transection | + | - | PRP | PRP alone | + | - | + | + |
| [55] | 2013 | Horse | SDFT | Collagenase gel | - | - | PC | Sham | + | - | - | - |
| [56] | 2014 | Horse | SDFT | Lesion created by a standardized surgical model | - | - | - | Sham | + | - | + | - |
| [57] | 2014 | Rabbit | CCT | Resection of a 2 cm-long tendon fragment | - | + | - | Cell-free scaffold | + | -- | - | + |
| [7] | 2014 | Rabbit | SST | Sharp release of the insertion of the SST at the greater tuberosity | +<sup>1</sup> | - | - | Sham | - | - | + | - |
| [58] | 2016 | Horse | SDF | Lesion created by a standardized surgical model | - | - | - | Sham | + | - | + | - |
| [59] | 2016 | Dog | FDPT | Transection | + | + | GF | Cell-free scaffold | - | - | - | + |
| [60] | 2016 | Dog | FT | Transection | + | + | - | Cell-free scaffold | - | - | + | - |
| [61] | 2017 | Horse | SDFT | Lesion created by a standardized surgical model | - | - | PRP | BM-MSCs | + | - | - | - |
| [62] | 2017 | Horse | SDFT | Lesion created by a standardized surgical model | - | - | - | Sham | + | - | - | + |
| [63] | 2017 | Dog | FDPT | Transection | + | + | GF | Cell-free scaffold | + | - | + | - |
| [64] | 2018 | Horse | SDFT | Collagenase | - | - | - | Autologous serum | + | + | + | - |
| [65] | 2019 | Rabbit | PT | Partial patellectomy | + | + | US | No cells, no US | + | +<sup>3</sup> | - | + |
| [66] | 2019 | Rabbit | FDST | Transection | + | - | - | Sham | - | - | - | + |
| [67] | 2019 | Sheep | AT | Collagenase | - | - | rESWT | PRP | + | - | + | - |
| [68] | 2020 | Rat | SST | Sharp release of the insertion of the SST at the greater tuberosity | +<sup>2</sup> | + | - | Cell-free scaffold | + | - | + | + |

Abbreviations: R, reference number; Y, year of publication; Sp, species; T, tendon; I, injury; P, primary repair; S, use of a scaffold; AT, additional treatment; C, control treatment; H, histology; PM, polarization microscopy; IHC, immunohistochemistry; B, biomechanical analysis; CCT, common calcaneal tendon; SDFT, superficial digital flexor tendon; SST, supraspinatus tendon; FDPT, flexor digitorum profundus tendon; FT, flexor tendon; PT, patella tendon; FDST, flexor digitorum superficialis tendon; AT, Achilles tendon; PRP, platelet rich plasma; PC, platelet concentrate; GF, growth factors; US, low-intensity pulsed ultrasound; rESWT, radial extracorporeal shock wave therapy. <sup>1</sup>three weeks after release of the insertion of the supraspinatus tendon at the greater tuberosity. <sup>2</sup>two weeks after release of the insertion of the supraspinatus tendon at the greater tuberosity. <sup>3</sup>polarization microscopy of the tendon-bone junction.

All of these studies reported benefits of treatment with cultured ADSCs. However, none of these studies are comparable with the present study. In particular, in the majority of these studies surgical transection of tendons (or sharp release of the tendon at the tendon-bone insertion site) was followed by primary repair, and cultured ADSCs were used to augment the surgical repair of the tendon. Furthermore, (i) in none of these studies detailed photomicrographs of histological sections demonstrating seamless integration of newly formed connective tissue with characteristics of tendon tissue into the existing tendon tissue were shown, (ii) none of these studies demonstrated that treatment with cultured ADSCs prevented the formation of blood vessels and adipose tissue at the junction between the original tendon tissue and the newly formed connective tissue, and (iii) none of these studies applied functional histology and functional immunohistochemistry as performed in the present study (c.f. Figs 16-20). In addition, in only one of these studies [65] polarization microscopy was used to demonstrate discernible crimp in the newly formed connective tissue (in [65] polarization microscopy was used to assess the tendon-bone junction). Moreover, none of these studies reported a substantial reduction in the relative amount of cells in the newly formed connective tissue over time after treatment with cultured ADSCs (c.f. Fig. 21).

Many recent studies addressed treatment of tendon injuries with allogeneic or xenogenic, cultured ADSCs in vivo (e.g., [69-71]). One potential motivation for the use of allogeneic cells in tendon regeneration could be the possibility to produce allogeneic, cultured ADSCs on an industrial scale without the need to perform on each individual subject a mini-liposuction before treatment [72]. An advantage of using allogeneic cells in treatment of tendon injuries would be to minimize biological variation [73], and allogeneic mesenchymal stem cells would provide the benefits of proven cellular purity, multipotency, consistency and ease of acquisition without the delays required for autologous cell isolation, characterization and propagation; these benefits would outweigh the risk of an inflammatory (rejection) reaction in response to foreign proteins, specifically major histocompatibility complexes [74]. On the other hand, the same study [74] found no superiority of treating an experimentally induced Achilles tendon defect in rats with allogeneic cultured ADSCs in fibrin gel over sham treatment with fibrin gel alone. This negative finding was in line with the finding that treatment of racehorse superficial digital flexor tendonitis with a combination of allogeneic cultured ADSCs and a controlled exercise rehabilitation program (CERP) was not superior to CERP alone [75], and a recent clinical, randomized controlled trial found no superiority of treating sPTRCT with allogeneic, cultured ADSCs over sham treatment with injection of saline [76]. In summary, these data challenge and contradict the supposed advantages of allogeneic cultured ADSCs in treatment of tendon pathologies outlined in [74].

One group proposed the use of allogeneic ADRCs (i.e., in general a similar approach to the one in the present investigation, but in an allogeneic manner) for treating tendon pathologies [77-79]. The corresponding studies were performed on a rabbit deep digital flexor tendon transection / primary repair model and demonstrated improvements in biomechanical properties of the tendons after augmenting primary repair with allogeneic ADRCs compared with primary repair sham-augmented with application of phosphate buffered saline [77-79]. On the other hand, histological and immunohistochemical analysis was only rudimentarily carried out in one of these studies [77], and the data shown in [71] do not allow to assess the quality of the newly formed tissue at the site of transection / primary repair. Moreover, it is reasonable to hypothesize that in the experimental defect / repair model investigated in [77-79] application of autologous UA-ADRCs would have led to better outcome than application of allogeneic ADRS. In this regard a study on experimental, orbital volume augmentation in rabbits with injection of hyaluronic acid and human, xenogenic ADRCs resulted in lasting inflammation for at least 6 weeks post-treatment followed by formation of profound fibrosis/scar tissue [80], which may have also occurred in [77-79]. However, formation of profound fibrosis/scar tissue (most probably secondary to a chronic inflammatory (rejection) reaction caused by application of xenogenic ADRCs) is neither the aim in orbital volume augmentation [81] nor the aim in tendon regeneration. Considering these data and substantial ethical concerns in potential application of allogeneic ADRCs in treatments of human tendon pathologies this approach appears not suitable for clinical routine use.

One limitation of the present study is that the exact molecular and cellular mechanisms of action of UA-ADRCs in tendon regeneration could not be determined, and will remain unknown. This is a likely consequence of the complex interactions between the different cell types contained in UA-ADRCs and their interactions with the host tissue that cannot be investigated either in vitro (as UA-ADRCs cannot be cultured in their entirety) or in vivo (as UA-ADRCs cannot be labeled when applied immediately after isolation, as done in the present stuy as well as in clinical applications [1,2,6,11,47]). Three examples of these complex interactions are briefly explained in the following (c.f. also [82]).

First, in tendon healing, stromal cell-derived factor-1α (SDF-1α) acts as chemoattractant, and transforming growth factor-β1 (TGF-β1), and insulin-like growth factor-1 (IGF-1) modulate cell proliferation and differentiation (reviewed in [83]). These growth factors are expressed in higher concentrations in uncultured ADRCs than in cultured ADSCs [83], and they can all be secreted by M2 macrophages [84-86]. Thus, the role of M2 macrophages contained in UA-ADRCs in treating tendon pathologies may go far beyond simple anti-inflammatory effects. Furthermore, mesenchymal stem/stromal cells (MSCs) contained in UA-ADRCs may suppress pro-inflammatory M1 macrophages [87] and convert M1 macrophages into M2 macrophages [88] in the target area. Second, the MSCs contained in UA-ADRCs can enhance the proliferation of tenocytes [87]. On the other hand, after injection of cultured autologous ADSCs into a rabbit Achilles tendon defect/repair model in vivo, the cells differentiated into tenocytes and integrated into the host tissue [24]. Furthermore, after injection of cultured autologous ADSCs into an experimentally induced tendon defect in horses in vivo, the ADSCs differentiated into cells that were integrated into new (tendon) tissue, with detection up to nine weeks post-treatment [58]. Moreover, when seeding human, cultured ADSCs on a specific scaffold (Hyalonect meshes) in vitro, the ADSCs created a capillary network within the scaffold [89]. These results indicate that after injection of UA-ADRCs into a tendon defect, the MSCs contained in the UA-ADRCs may differentiate into other cell types that are necessary for tendon regeneration and integrate into the host tissue. However, these results also imply that in the model investigated in the present study one cannot discern whether the cells found in the newly formed connective tissue arose by proliferation of local cells or by proliferation and differentiation of cells contained in UA-ADRCs.

Third, human type I collagen expression was found in rat Achilles tendon tissue four weeks after treating an experimental, full-thickness, intratendinous, rectangular Achilles tendon defect (0.8 mm x 5 mm) by application of human, cultured ADSCs [90], which cannot be attributed to enhanced production of type I procollagen by host (rat) tendon cells. Thus, the type I procollagen observed in the present study in newly formed connective tissue after application of UA-ADRCs may have been produced by cells contained in the UA-ADRCs and/or by local cells.

From a clinical perspective, elucidating the exact mechanisms of action of UA-ADRCs in tendon regeneration is of secondary importance: any selection, cultivation, stimulation, manipulation and/or (genetically) reprogramming of cells contained in ADRCs aiming at optimizing their mechanisms of action in tissue regeneration is incompatible with meeting the criteria of ‘minimally manipulated’ as defined in 21 CFR 1271.10(a) [91] (Title 21 is the portion of the Code of Federal Regulations that governs food and drugs within the U.S. for the FDA [92]), preventing regulation solely under Section 361 of the Public Health Service (PHS) Act and 21 CFR Part 1271 [93]. The European Medicines Agency considers such “optimized” cells as an Advanced Therapy Medicinal Product (ATMP) [94], which is not the case for human UA-ADRCs isolated from adipose tissue with the technology used in the present study. Rather, from a clinical perspective the composition of UA-ADRCs is of primary importance, and high relative amounts of MSCs, endothelial progenitor cells and M2 macrophages are most desired. Of all systems and technologies available for isolating ADRC from adipose tissue, the technology applied in the present study results in the highest relative numbers of these cell types [5]. Furthermore, the relative numbers of these cell types as well as other key characteristics of UA-ADRCs (number of nucleated cells, cell viability and number of viable nucleated cells per gram of adipose tissue harvested) were independent of the subject’s age, sex, body mass index and ethnicity [5].

Another limitation of this study is that an acute tendon defect model was used, whereas the vast majority of partial-thickness rotator cuff tears are not acute but develop in tendons which present the failed healing lesion typical of tendinopathy [95-97]. On the other hand, the perfect animal model for tendinopathy does not exist, and models should be carefully considered in light of the specific research question [98]. In line with this the animal model used in the present study would be inadequate for investigating the efficacy of orthobiologics that aim at stimulating tissue-resident cells such as platelet rich plasma (PRP) [99-101] or exosomes derived from cultured stem cells [102-104]. In this regard, a recent double-blinded, randomized controlled clinical trial concluded that injections of PRP might not be beneficial in non-operative treatment of rotator cuff disease [105,106]. In contrast to PRP and exosomes the application of UA-ADRCs primarily focuses on pathologies in which the patient’s body’s localized self-healing power is exhausted and, as a consequence, physiological body structures and functions can no longer be restored by the local stem cell pool [23]. The tendon defect model used in the present study is adequate for this purpose.

Finally we would like to emphasize that for the understanding of tissue regeneration or regenerative medicine in the context of clinical application the type of animal study presented here is perhaps even more significant than in treating other conditions. The implications here are that decreased pain and increased mobility would result from this type of treatment in humans. However, decreased pain and increased mobility are much more subjective clinical endpoints than in many therapeutic areas. Clearly, imaging rather than examination of biopsies will have to be relied upon in a clinical setting to determine whether regeneration occurred. Thus, the present study shows that tissue regeneration that clearly would not be an ascertainable measure in humans has been demonstrated, and treatment of tendon pathologies with UA-ADRCs has the potential to be truly “structure-modifying” rather than just being “symptom-modifying” [107].

## DECLARATIONS

### Ethics approval

The animal study protocol was approved by the Colorado State University (CSU) Institutional Animal Care and Use Committee (Fort Collins, CO, USA) (Protocol # 1473; approval issued on February 1st, 2021 and renewed/amended on February 1st, 2022).

### Consent for publication

Not applicable

### Availability of data and materials

The datasets used and analyzed during the current study are available from the corresponding author on reasonable request.

### Competing interests

C.S. is Advisory Medical Director of InGeneron, Inc. (Houston, TX, USA). C.A. is Director of Medical and Scientific Affairs of InGeneron. E.A. is Executive Chair and President of InGeneron. However, InGeneron had no role in study design, data collection, and analysis, interpretation of the data, and no role in the decision to publish and write this manuscript. No other potential conflicts of interest relevant to this article were reported.

### Funding

This research received no external funding.

### Authors’ contributions

CS has made substantial contributions to the conception and the design of the work, to the acquisition, analysis and interpretation of data, and has drafted the work. CA has made substantial contributions to the conception and the design of the work, to the analysis and interpretation of data, and has substantially revised the work. TW and SM have made substantial contributions to the acquisition, analysis and interpretation of data, and have substantially revised the work. JD and AH have made substantial contributions to the acquisition of data. KS, LB, JE, HS, CP, BG and KL have made substantial contributions to the conception and the design of the work as well as to the acquisition, analysis and interpretation of data, and have substantially revised the work. DP and NM have substantially revised the work. EA has made substantial contributions to the conception and the design of the work as well as to the analysis and interpretation of data, and has substantially revised the work.

All authors have approved the submitted version, and have agreed to be personally accountable for the author’s own contributions and to ensure that questions related to the accuracy or integrity of any part of the work, even ones in which the author was not personally involved, are appropriately investigated, resolved, and the resolution documented in the literature.

## Acknowledgments

The authors thank Beate Aschauer, Andrea Haderer and Claudia Harbauer for skillful technical assistance.

